# Chemical signals act as the main reproductive barrier between sister and mimetic *Heliconius* butterflies

**DOI:** 10.1101/856393

**Authors:** M.F. González-Rojas, K. Darragh, J Robles, M. Linares, S Schulz, W.O McMillan, C.D Jiggins, C Pardo-Diaz, C Salazar

**Affiliations:** Department of Biology, Faculty of Natural Sciences and Mathematics, Universidad del Rosario, Bogota, Colombia, 111221; Department of Zoology, University of Cambridge, Cambridge, Cambridgeshire, United Kingdom, CB2 3EJ; Department of Chemistry, Pontificia Universidad Javeriana, Bogota, Colombia; Institute of Organic Chemistry, Technische Universität Braunschweig, Germany; Smithsonian Tropical Research Institute, Panama

**Keywords:** *Heliconius*, Pheromone, Mate Choice, Speciation, Reproductive Isolation

## Abstract

Colour pattern has been long recognised as the trait that drives mate recognition between *Heliconius* species that are phylogenetically close. However, when this cue is compromised such as in cases of mimetic, sympatric and closely related species, alternative mating signals must evolve to ensure reproductive isolation and species integrity. The closely related species *Heliconius melpomene malleti* and *H. timareta florencia*, occur in the same geographic region and despite being co-mimics they display strong reproductive isolation. In order to test which cues differ between species, and therefore potentially contribute to reproductive isolation, we quantified differences in wing phenotype and male chemical profile. As expected, wing colour pattern was indistinguishable between the two species while the chemical profile of their male sex pheromones showed marked differences. We then conducted behavioural experiments to study the importance of these signals in mate recognition by females. In agreement with our previous results, we found that pheromones and not wing colour pattern drive the preference of females by conspecific males. In addition, experiments with hybrid males and females suggested an important genetic component for both pheromone production and preference. Altogether, these results suggest that pheromones are the major reproductive barrier opposing gene flow between these two sister and co-mimic species.

## INTRODUCTION

The mechanisms by which species maintain their integrity are diverse and usually involve the combination of multiple signals of intra and inter-specific communication such as chemical, visual, auditory and tactile cues (1–5). In particular, sexual communication in insects involves long and short-range pheromones, which play multiple roles (3,4,6,7). For instance, pheromones inform the mating status of females (8,9), quality and age of males (10,11), quality and quantity of nuptial gifts (12), body size (13), dominance status (14), and degree of relatedness (15). Also, sex pheromones play an important role in mate choice and species recognition (16,17). In particular, pheromones mediate mate choice in many insects including flies (*Drosophila*), grasshopers (*Chorthippus parallelus*), leaf beetles (*Chrisochus*) and stick insects (*Timema*) (16–32).

In Lepidoptera, both males and females produce volatile and non-volatile compounds suggesting that chemical communication plays a critical role in inter and intra-specific communication (33–35). In *Bicyclus anynana*, visual and chemical cues are equally important for mate choice, and females recognize heterospecific males based on their pheromones (36). Also, males of *Heliconius charithonia* which are known to engage in pupal mating by copulating with females as they eclose, are able to identify the sex of a conspecific pupa based on sex-specific compounds (37,38). Moreover, sex pheromones seem to mediate species recognition among mimetic and distantly related *Heliconius* where conflicts between mimicry and sexual communication may arise (39–41). This is supported by the fact that males of *Heliconius* have species-specific mixtures in their wing androconia (41,42).

In *Heliconius*, mate discrimination between closely related species relies on wing colour pattern (43,44). For example, the sister species *H. melpomene* and *H. cydno* (divergence ~1.5–2 Mya) (45), which are sympatric across Central America and the Andes, not only differ in habitat use but also in wing colouration (46–48). In fact, multiple experiments show that males prefer to court females exhibiting their own colour pattern (43,49,50). In contrast, the phylogenetically close *H. melpomene malleti* and *H. timareta florencia*, mimic each other and coexists in sympatry in the South Eastern Andes of Colombia (Figure S1). Despite their phenotypic resemblance, this species pair shows strong pre-mating ecological isolation (differences in host plant preference) as well as strong reproductive isolation tested in no-choice experiments (49,51); even so, few hybrids are found in nature (~2%) (40,49). Therefore, the strong reproductive isolation observed between *H. m. malleti* and *H. t. florencia* implies that sexual isolation is mediated by cues other than colour pattern, such as sexual pheromones (40,41,50).

In agreement with this hypothesis, previous studies showed that the two species differ in their androconia and genital chemical composition, although these studies included few individuals (39,41). Furthermore, a previous study of ours showed that females of *H. m. malleti* and *H. t. florencia* strongly discriminated against conspecific males which have their androconia experimentally blocked, affecting reproductive success with implications for reproductive isolation. This suggests that chemical signalling is important in mate choice in *Heliconius* butterflies (52).

Nonetheless, the importance of sex pheromones in mate recognition in *Heliconius* still needs further investigation. Specifically, we need to investigate more on: (i) the preference of females for conspecific vs. heterospecific male pheromones, and (ii) the inheritance patterns of both male pheromone production and female preference for them. Here, we used a combination of behavioural and chemical analyses to get a better understanding of reproductive isolation meditated by chemical signals in *Heliconius* butterflies.

## METHODS

### Wild sampling and interspecific crosses

For the analysis of colour quantification, we used wings of wild males of *H. t. florencia* and *H. m. malleti* deposited in the ‘Colección de Artrópodos de la Universidad del Rosario (CAUR229)’ (Table S1). We also collected wild individuals of *H. t. florencia* and *H. m. malleti* in the localities Sucre and Doraditas in Colombia (01°48’12" N −75°39’19" W, 1200 m and 01°42’39" N - 75°42’32" W, 1400 m) that were taken to the insectaries of the Universidad del Rosario in La Vega (Colombia) in order to establish stock populations for the behavioural experiments and chemical analyses. Larvae were reared on *Passiflora oerstedii* while adults were provided with *Psiguria* sp. as pollen source and supplied with ~20% sugar solution. We also used the stock populations to perform interspecific crosses between *H. t. florencia* and *H. m. malleti.* To do so, a female of *H. t. florencia* was mated with a male of *H. m. malleti* (the reciprocal cross was successful three times but the female always died before laying eggs). Then, two F_1_ males were backcrossed (BC) to pure *H. t. florencia* females (crosses towards females of *H. m. malleti* consistently failed). In all cases, eggs were collected daily and placed in small plastic pots. Larvae were reared individually to avoid cannibalism, and right before pupation, they were transferred to bigger plastic pots until eclosion. The two backcrosses towards *H. t. florencia* produced 25 males and 25 females: all of these males were processed to chemically characterise the composition of sex pheromones, while 24 females were used in behavioural experiments testing for female preference (see below).

### Quantification of wing phenotype

We evaluated whether *H. m. malleti* and *H. t. florencia* exhibit differences in wing phenotype. In order to do this, we scanned ventral and dorsal forewings (FW) and hindwings (HW) of 43 *H. m. malleti* and 45 *H. t. florencia* (Table S1) using a high-resolution flatbed scanner Epson Perfection V550, in RGB colour format with a resolution of 2400 dpi. Right side wings were always used. Then, we used ImageJ (53) to place a set of 34 landmark coordinates (LM) per individual (18 on the FW and 16 on the HW, dorsally and ventrally; Figure S2A). These LM were analysed in the R package Patternize (54) to quantify variation in wing colour patterns (band size and shape). This package extracts, transforms and superimposes colour patterns to finely quantify the variation of colour pattern phenotypes among species and performs a principal component analysis (PCA) (54). We then tested differences in wing pattern among species using a multivariate analysis of variance (MANOVA) in R based on a subset of PCs (those that explain >95% of the variation).

In addition, we used tpsDig2 (55) to place 32 landmark coordinates on the outline of both FW and HW (dorsally; Figure S2B). These LM coordinates were superimposed using a General Procrustes Analysis (GPA) in the R package ‘geomorph’ (56–58). GPA aligns, scales and rotates the configurations to line up the corresponding landmarks as closely as possible, minimizing differences between landmark configurations. The resulting coordinates in the tangent space were used as shape data, while the log-transformed centroid size (CS, hereafter) (57) was used as a size estimate (59). Differences in wing size among species were investigated with a one-way ANOVA with size as a dependent variable and species as a factor, followed by Tukey’s pairwise comparison test. The differences were visualised with a boxplot. Differences in wing shape among species were tested using a Procrustes MANOVA applied to the aligned landmark configurations. This was done using the *procD.lm* function in the ‘geomorph’ R package (56).

### Behavioural experiments

In order to test female preference for colour pattern and sexual pheromones of conspecific males, we conducted two types of behavioural experiments in triads: (i) altering the wing phenotype of males, and (ii) perfuming males with the heterospecific pheromone mixture. All experiments were conducted from 7 am to 1 pm, checking every 30 minutes for mating; the experiments stopped when mating occurred. For each experiment, mature males (at least 10 days old) were randomly selected from the stock population while females were used as soon as they became available. If no mating occurred on the first day, we repeated the experiment the next day using the same butterflies; in contrast, mated males or females were never reused. If no mating occurred after the second day, the experiment stopped. For the first hour of each experiment, we observed female behaviour towards the males. These behaviours were recorded only when a male was actively courting the female. Observations were divided into one-minute intervals and we recorded female behaviours of “acceptance” or “rejection” previously defined for *Heliconius* (52,60). Specifically, we recorded the following acceptance behaviours: flutter (female does a slow and moderate wing flapping), fly towards (female flies facing towards the male), slow flat (female does a slow rhythmic flight), and wings open and exposed (female opens her wings and her abdomen is relaxed). Similarly, we recorded the following rejection behaviours: fly away (female flies away from the male), tucked up (female is alighted with her wings closed and her abdomen concealed within the wings), rapid and erratic flutter (female does a high frequency flutter of her wings and raises her abdomen when the male is close proximity), and wings opened and abdomen bend (female has her wings opened and her abdomen raised, but without wing fluttering).

In the triads that tested female preference for colour pattern, a single one-day old virgin female was presented with two conspecific males of at least ten days old (i.e. sexually mature). One of the males (treatment) had his FW and HW completely blacked-out using a black marker (COPIC 100) thus hiding his wing colour pattern. The second male (control) had his FW and HW painted with a colourless marker (COPIC 0); in this way, the male kept his phenotype unaltered, but we controlled for any odour effect of the marker. We tested a total of 20 females per species.

For the triads that tested female preference for sexual pheromones, we first prepared pheromone extracts from sexually mature males of each species by dissecting and mixing the androconia region of five conspecific individuals of the same age and soaking them in 200 µL of hexane for 1 hour. After this incubation, the solvent was transferred to a new vial and stored at −20°C until needed. Then, a single one-day old virgin female was presented with two sexually mature conspecific males. Both males were initially treated with transparent nail varnish applied on the dorsal side of their HW in order to block their androconia and thus, block their natural pheromone emission. Then, one of the males (control) was perfumed by spreading the conspecific hexane extract in the androconia region, whereas the second male (treatment) was perfumed by spreading the heterospecific hexane extract. A total of 19 females of *H. t. florencia* and 18 females of *H. m. malleti* were tested. In order to investigate how long the hexane extract remains in the wings of the perfumed males before completely evaporating (which can potentially affect our results), we blocked the HW androconia of nine males of each species using transparent nail varnish and then, we re-perfumed this region by spreading the heterospecific hexane extract. We left these males fly in an insectary and dissected their androconia at 1 minute, 30 minutes or 60 minutes and soaked this tissue in 200 µL of ultrapure dichloromethane (Merck UniSolv®) to be later analysed by gas chromatography/mass spectrometry (GC/MS; see below).

In addition, we studied hybrid female preference. A single virgin hybrid female had to choose between two males, one *H. t. florencia* and one *H. m. malleti.* A total of 18 F_1_ and 24 BC one-day old virgin females were tested. These triads were conducted in the same way as the previous behavioural experiments. Female acceptance or rejection behaviours were also recorded.

The mating outcome was analysed with a binomial test. We also used a generalised linear mixed model (GLMM) with a binomial error distribution and logit link function to test if females responded differently to control and treatment males. The response variable was derived from those minutes where at least one of the males courted the female regardless of her response (either “acceptance” or “rejection”). Significance was determined by using likelihood ratio tests comparing models with and without male type included as an explanatory variable. In order to avoid pseudoreplication, individual female was included as a random effect in all models. All statistical analyses were performed with R version 3.3.2 (61), using the packages ggplot2 (62), car (63) and binom (64) following Darragh et al. 2017 (52).

### Characterisation of chemical profiles

In order to determine the chemical composition of androconia wing tissue and genitalia bouquet in males of *H. m. malleti, H. t. florencia* and their hybrids (F_1_ and BC), we dissected both tissues from adults and placed them individually in 200 µL of ultrapure dichloromethane (Merck UniSolv®) in 2 mL glass vials and soaked for 1 hour. After this incubation, the solvent was transferred to a new vial and stored at −20°C. Dichloromethane is a preferred solvent used for pheromone extraction since it is sufficiently volatile for extracts to be concentrated without exposing them to high temperatures, it is non-flammable, and penetrates scales better than hexane leading to higher pheromone titles (65,66).

These extracts were analysed by gas chromatography/mass spectrometry (GC/MS) at the Smithsonian Tropical Research Institute in Panama following the protocol of Mann et al., 2017 (41). Prior to the GC/MS, samples were evaporated under ambient air at room temperature. We quantified the compounds found in the extracts using GC with flame ionisation detection with a Hewlett-Packard GC model 7890B equipped with a Hewlett-Packard ALS 7693 autosampler. A BPX-5 fused silica capillary column (SGE, 25m × 0.22 mm, 0,25 µm) was used. Injection was performed in splitless mode (250°C injector temperature) with hydrogen as the carrier gas with a constant flow of 1.65 ml/min. The temperature programme started at 50°C, held for 5 min, and then rose to 320°C with a heating rate of 5°C/min. We used 2-tetradecyl acetate (200 ng) as an internal standard. Components were identified by comparison of mass spectra and gas chromatographic retention index with reference samples from the Schulz lab collection (Institute of Organic Chemistry, Technische Universität Braunschweig, Germany). Relative concentrations were determined by peak area analysis by GC/MS.

Species differences in compound composition were evaluated with a canonical discriminant analysis applied on PCAs accounting to 95% of the variance. Dimension reduction (PCAs) was implemented in the software Past v3.0 (67). A MANOVA was performed in candisc R package (68). Finally, using the means of the relative concentrations of each compound, we established the relationship among *H. t. florencia*, *H. m. malleti*, F_1_s and BCs males for androconia and genital bouquet, calculating dendrograms using Euclidian distance. Both, dendrogram and compound composition were visualised using the function heatmap in R (version 3.5.0). GC/MS raw data is available in the public repository Zenodo (https://zenodo.org/).

## RESULTS

### Quantification of wing phenotype

We found that *H. m. malleti* and *H. t. florencia* are significantly different in wing size, both in the FW and HW (ANOVA p=0.00001 in both cases, Figure S3A and 3B). FW and HW size in *H. t. florencia* is consistently larger than in *H. m. malleti*. In terms of shape, the FW is statistically different between species (MANOVA p=2.504e-12), but not the HW (MANOVA p=0.0371; α=0.01, Figures S3C, S3D and S4A). In the FW, the most variable LMs were LM3, LM4 (located in the distal part of the costal margin, near the wing apex; Figures S2C, S2D and S4B) and LM15 (located in the middle of the inner margin; Figures S2E, S2F and S4C). Thus, the significant shape variation between the two species appears to be linked with the length of the FW (longer in *H. t. florencia*) and the curvature of the inner margin (deeper in *H. t. florencia*).

The colour pattern comparison between the two species suggested subtle differences in wing colour pattern (Figure S5). The shape (PC2) of the three wing elements investigated (forewing band, forewing ‘dennis’ patch, and hindwing rays) did not differ between species, but their size (PC1) was slightly different (Figure S5). The edge of the yellow forewing band seemed more distally extended in *H. m. malleti* but only on the ventral side of the wing (Figure S5A, S5D). Also, the hindwing rays appeared thicker in *H. m. malleti*, but only on the dorsal side of the wing (Figure S5C, S5F). In contrast, the ‘dennis’ forewing patch is bigger in *H. m. malleti*, both dorsally and ventrally (Figure S5B, S5E). Despite this, none of these differences were statistically different between species indicating that the two species are almost indistinguishable in terms of wing colour pattern.

### Behavioural experiments

Altering the wing pattern of males had no effect on mating probability. In all 40 experiments, both *H. t. florencia* and *H. m. malleti* females mated readily with conspecific males completely blacked (p=0.55 in both cases; Figure 1A). Consistent with this finding, we found no differences in female response towards control and treatment males in all but one of the acceptance and rejection behaviours we assayed (Figure 1B and Table S2). The exception was the observation that female *H. t. florencia* kept their wings open for a larger proportion of the time during courtship by control males relative to male whose wing patterns were eliminated (open wings; Figure 1B).

**Figure 1.**
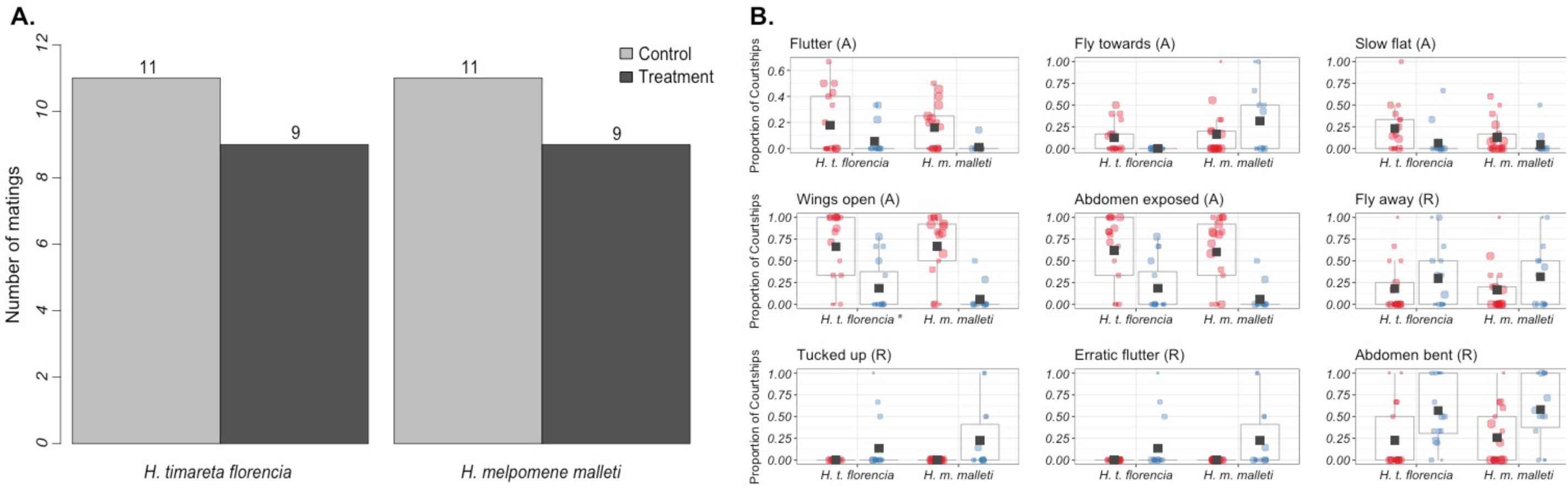
(**A**) Mate choice triads testing the importance of wing colour pattern in mate preference. Number of matings obtained is indicated above each bar. Control males are represented in light grey and treatment males are represented in dark grey. (**B**) Proportion of courtships, which resulted in female behavioural responses (Acceptance (A) or Rejection (R)) towards control and treatment males. Control males are represented in red (left) and treatment males in blue (right). Means are marked with a black square and boxplots mark the inter-quartile ranges. Size of datapoint is proportional to the number of courtships by that male. Asterisk (*) next to the species’ name is indicative of statistically significance (α=0.01) according to GLMM.

The triads (n= 37) testing female preference for perfumed males did not yield any matings. The failure to mate might reflect the rapid evaporation rate of the chemical extracts. For example, the concentration of octadecanal decreased 25% in the first 30 minutes and 82% after 60 minutes (Figure S6 and Table S4). Similarly, the concentration of syringaldehide decreased 25% in the first 30 minutes and 47% after the first hour. Nevertheless, we observed significant differences in female behavioural responses towards treatment and control males (Figure S7 and Table S3). Specifically, in all behaviours tested females exhibited acceptance behaviours towards males perfumed with the hexane extract of their own species and consistently rejected males perfumed with that of the other species.

F_1_ and BC females obtained from mating an F_1_ female to a *H. t. florencia* male were reluctant to mate either parental species. In these trials, F_1_ mated at 33% frequency and BC females at 37% frequency; in all cases matings were with *H. t. florencia* males (Figure S8). This agrees with previously unpublished no-choice experiments where F_1_ mated at 25% frequency and BC females at 37% frequency, always with *H. t. florencia* males (Table S8). Consistently, hybrid females were more likely to perform acceptance behaviours towards *H. t. florencia* males while rejection behaviours were observed more often towards *H. m. malleti* males (Table S5).

### Characterisation of chemical profiles

We analysed a total of 100 wing androconia and 95 abdominal gland extracts from males of *H. m. malleti*, *H. t. florencia*, F_1_ and backcrosses to *H. t. florencia*. Males of both species presented a common composition in the wing androconia extract, and the most notable differences were in terms of the concentration of individual compounds in the blend (Table S6). The compounds of the androconial region in *H. m. malleti* were mainly alkanes (28%), aldehydes (16.27%) and unknown compounds (18%), with octadecanal being the most abundant compound (1094.91 ng in *H. m. malleti* compared to 1.40 ng in *H. t. florencia*). In *H. t. florencia*, the androconial bouquet was composed of alkanes (37%) and esters (14%), with syringaldehide and heneicosane being the most abundant compounds (Figure S9). The androconial composition of F_1_ and BC males were very similar to that of *H. t. florencia* but showed higher individual variation (Figure S10, Figure S11; Table S6). Consistently, the discriminant analysis revealed a discrete group formed by *H. t. florencia*, F_1_ and BC males, while *H. m. malleti* formed an independent cluster (Figure 2A; MANOVA p<2e-16). In addition, a post-hoc Tukey test showed that F_1_ and BC males differed significantly from *H. m. malleti* males (p=0.00000001; in both cases) but not from those of *H. t. florencia* (p=0.912036 and p=0.8915162; respectively).

**Figure 2.**
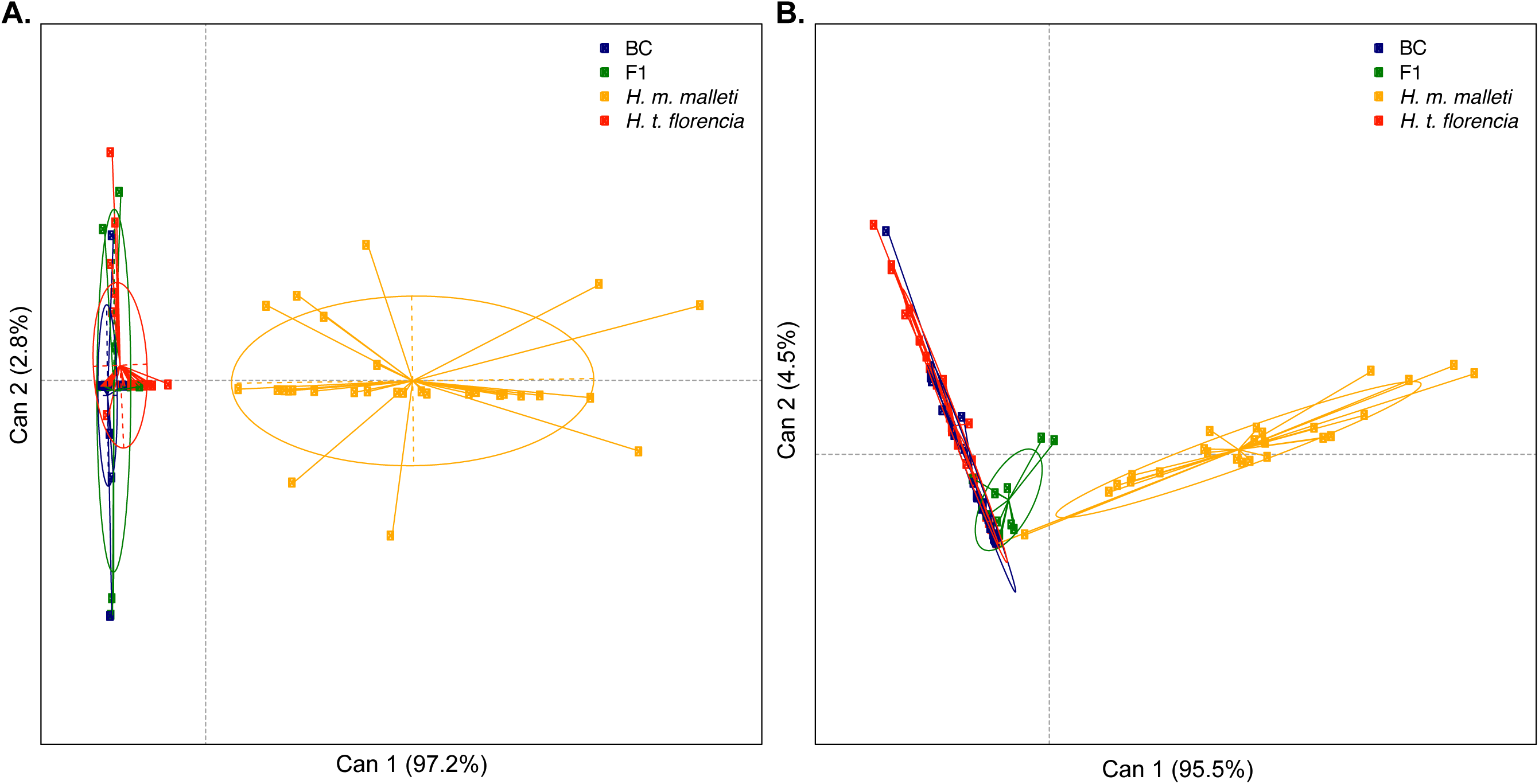
(**A**) Discriminant analysis based on individual composition of wing androconia extracts of males of *H. m. malleti*, *H. t. florencia*, F_1_ and BC. (**B**) Discriminant analysis based on individual composition of abdominal gland extracts of males of *H. m. malleti*, *H. t. florencia*, F_1_ and BC.

The abdominal gland bouquet of males was chemically more diverse than that of the androconia (Table S7). Alkanes, esters and lactones were the major compounds present in the abdominal gland bouquet of *H. m. malleti* and *H. t. florencia*. β-Ocimene and heneicosane largely dominated the abdominal gland bouquet of *H. m. malleti* males, while that of *H. t. florencia* was mainly composed of ethyl oleate, butyl oleate, isopropyl oleate and (Z)-9-octadecen-13-olide (Table S7, Figure S12). As in the androconia, the abdominal gland bouquet of F_1_ and BC males was more similar to *H. t. florencia* although some individuals had β-ocimene (Figure S13 and Figure S14). The discriminant analysis revealed a discrete group composed of *H. t. florencia*, F_1_ and BC males which differentiate from a cluster formed only by *H. m. malleti* (Figure 2B; MANOVA p<2e-16). A post-hoc Tukey test showed that F_1_ and BC males differed significantly from *H. m. malleti* males (p=0.00000001; in both cases) but not from *H. t. florencia* males (p=0.1521143 and p=0.7778819; respectively). Interestingly, both species and their hybrids showed putative defensive secretions in their abdominal gland, namely 2-sec-butyl-3-methoxypyrazine in the abdominal gland extracts of *H. t. florencia*, F_1_ and BC and 2-isobutyl-3-methoxypyrazine in the abdominal gland extracts of *H. m. malleti*, F_1_ and BC (Table S7 and Figure S13). These specific compounds are known deter predators in the wood tiger moth (69), and in general, methoxypyrazines are compounds frequently found in the chemical defences of aposematic insects (70–72).

## DISCUSSION

In *Heliconius* butterflies, reproductive and behavioural isolation usually follow a multimodal pattern where multiple traits such as wing colouration, pheromones, and habitat use, among others, are involved (40). For example, visual cues mediate the recognition of conspecific females by males across the entire spectrum of divergence in the *melpomene/cydno/timareta* clade with implications for reproductive isolation and speciation (40). Thus, reproductive isolation between taxa with shifts in wing colour pattern, such as the species *H. m. rosina* and *H. c. chioneus* (high divergence) or the subspecies *H. t. linaresi* and *H. t. florencia* (low divergence), heavily relies on visual cues (40). In addition to wing colouration, chemicals seem to also contribute to species recognition (39,52,73). In fact, species identity largely explains the overall variation in chemical profiles in *Heliconius* (42).

To date, the contribution of wing colouration to reproductive isolation on *Heliconius* has been deeply studied and experimentally tested, mainly in males (43,44,51,74–77; although see 78). However, the role of pheromones and female preference in maintaining species integrity is far less understood. The convergence in wing colouration between the sister and sympatric species *H. t. florencia* and *H. m. malleti* provides a natural experimental system to: (i) disentangle the effect of wing colour pattern and chemical signals, and (ii) characterise the role of female preference in reproductive isolation. In this study we confirmed the existence of strong mating preferences between *H. m. malleti* and *H. t. florencia*, and this preference appears to not be related to colour pattern, but chemically mediated.

We found that *H. t. florencia* and *H. m. malleti* are nearly identical in terms of wing patterning, and in consequence, altering this trait with black markers had little effect on the preference of females for mates. This also accords with our earlier observations where experiments with wing models washed in hexane revealed that males of *H. t. florencia* approached and courted models of *H. m. malleti* as much as theirs (51). These results suggest that wing phenotype is not the cue that maintains species integrity between these mimetic pair. Instead, strong mating isolation appears to be largely driven by chemical signals. Our previous work demonstrated that females of *H. m. malleti* and *H. t. florencia* strongly discriminated against conspecific males that have their wing androconia experimentally blocked (52). Consistent with this result, we found that females showed more acceptance behaviour and less rejection behaviour towards males perfumed with a conspecific extract relative to those perfumed with the heterospecific extract. Moreover, the androconia scale cells of the two species had highly divergent profiles composed of differences in both the relative abundance and presence/absence of volatile chemicals. Therefore, this study along with previous findings show that assortative mating in *Heliconius* involves chemical signals (39,52). In cases of sister species that differ in wing colour pattern, both wing colouration and chemical cues have a combined contribution to isolation (79); as such, natural hybrids between the phenotypically divergent *H. m. rosina* and *H. c. chioneus* are rare (0.05%; 40). In contrast, in cases of mimicry between sister species where divergence in colour pattern was lost due to introgression (80,81) such as *H. m. malleti* and *H. t. florencia*, the occurrence of natural hybrids is more frequent (2-3%; 49). Therefore, our study is one of the few that provides experimental evidence that reinforces the importance of chemicals and female preference in incipient behavioural isolation in *Heliconius*.

In agreement with this hypothesis, we also found that the composition of the male pheromone bouquet of *H. m. malleti* and *H. t. florencia* is different, confirming the results previously reported in other studies that included a much lower sample size (39). The pheromone signature of the two species is unique, not only in the androconia but also in the genitalia. The pheromone mixtures differed between species mainly in the concentration of compounds and not in the composition, although there were some species-specific compounds (less abundant) (42). In particular, we identified octadecanal and β-ocimene as the main compounds in the androconia and genitalia bouquet of *H. m. malleti*, respectively. Even though octadecanal was also present in *H. t. florencia*, its abundance was much lower in this species. This agrees with recent findings, where octadecanal was found to be abundant in males of *H. m. rosina* and almost absent in those of *H. c. chioneus* (79). Interestingly, octadecanal has been proven to be electro-physiologically active in *Heliconius* (79) and β-ocimene is a known anti-aphrodisiac in the genus (82). On the other hand, the most abundant compounds in the male androconia of *H. t. florencia* were syringaldehyde and heneicosane, known to act a long-range attraction molecules in multiple insect species (83–87). Similarly, the male pheromone of this species also contained phenylacetaldehyde and limonene, which have been previously reported as copulating pheromones, hormones and defensive secretions in other Lepidoptera (84).

Hybrid males (F_1_ and BC) had fewer and less abundant compounds in their blends compared with the parental species. Interestingly, both F_1_s and BCs had little octadecanal and some hybrid individuals even lacked it at all (just as pure *H. t. florencia* males do) suggesting that the low amount of this compound is heritable and possibly involving few major effect loci. This agrees with recent findings in *H. m. rosina* and *H. c. chioneus* that suggest a potential monogenic basis and dominant inheritance for octadecanal production (79). Similarly, hybrid females (F_1_ and BC) accepted better males of *H. t. florencia* rather than those of *H. m. malleti*, and in some cases, we observed matings with the former species. This again suggests that female preference could be heritable and with a simple genetic basis. However, our data is not enough to draw definitive conclusions on the genetics of pheromone production or female preference. To answer this question, it would be necessary to test the reciprocal F_1_ (*H. m. malleti* mother x *H. t. florencia* father) and backcrosses towards *H. m. malleti*. However, such crosses are extremely difficult to obtain, perhaps due to intrinsic behavioural sterility as reported for other Lepidoptera (88).

Overall this study corroborates the existence of chemical differences between male sex pheromones in a pair of closely related and mimetic species where wing phenotype is not a recognition trait. These chemical differences are used by females to effectively choose mates. These results experimentally confirm that male sex pheromones are necessary to mediate premating reproductive isolation between *H. m. malleti* and *H. t. florencia*. Also, our results suggest that both male pheromone production and female preference in *Heliconius* are inheritable traits. In fact, recent studies have pinpointed candidate regions implicated in the production of male pheromones in *Heliconius* (79). However, to date, the genetic basis controlling female assortative mating behaviours remain unknown. In *Heliconius*, females exert choice based on multimodal signals, especially visual and olfactive cues (78). Therefore, in the absence of wing colour pattern divergence, the mimetic pair composed by *H. m. malleti* and *H. t. florencia* provides an excellent natural system to investigate the genetics of female preference based on pheromones and understand how species integrity is maintained in the presence of gene flow. It is possible that sexual signals and responses to those signals involve the same gene, such as in the case of *Drosophila melanogaster* (89), or that both traits are controlled by independent loci, such as in *Heliothis* (90).

## ACKNOWLEDGMENTS

We thank our team at the insectaries in La Vega (Colombia), including Oscar Penagos and Sebastian Sánchez for helping rearing the crosses used in this study. Steven Van Belleghem helped running Patternize, and Kelsey Bryes gave useful feedback. This work made use of the CENTAURO Supercomputer of the Universidad del Rosario High Performance Computing Service. This project was funded by the Universidad del Rosario FIUR Grant QFN-DG001 and COLCIENCIAS (Grant FP44842-5-2017). We also thank the Autoridad Nacional de Licencias Ambientales (ANLA) in Colombia for granting us the collection permit 530. SS thanks the Deutsche Forschungsgemeinschaft (DFG) for support through grant Schu984/12-1. CDJ was funded by a European Research Council grant number 339873 Speciation Genetics.

