## Supplemental Material for "Chemical signals act as the main reproductive barrier between sister and mimetic *Heliconius* butterflies"

Table S1. Samples included in the quantification of wing phenotype analyses. A total of 89 individuals were used. The wings were obtained from “Colección de Artrópodos de la Universidad del Rosario (CAUR229)”. Last column indicates analysis in which that specimen was used, D, dorsal; V, ventral; HW, hindwing; FW, forewing.

| **ID Collection** | **ID Wing Scan** | **Taxon** | **Locality** | **Analysis** |
| --- | --- | --- | --- | --- |
| M54 | LGE-WS-00351 | *H. t. florencia* | Quebrada_Las_Doraditas | D-HW; D-FW |
| M63 | LGE-WS-00349 | *H. t. florencia* | Finca_Piñacue | D-HW; D-FW; V-HW; V-FW |
| M64 | LGE-WS-00350 | *H. t. florencia* | Finca_Piñacue | D-HW; D-FW; V-HW; V-FW |
| M244 | LGE-WS-00373 | *H. m. malleti* | Florencia | D-HW; D-FW; V-HW; V-FW |
| M253 | LGE-WS-00375 | *H. m. malleti* | Florencia_Sucre | D-HW; D-FW; V-HW; V-FW |
| M255 | LGE-WS-00326 | *H. t. florencia* | Florencia_Sucre | D-HW; D-FW |
| M257 | LGE-WS-00346 | *H. t. florencia* | Florencia_Sucre | D-HW; D-FW; V-HW; V-FW |
| M259 | LGE-WS-00325 | *H. t. florencia* | Florencia_Sucre | D-HW; D-FW |
| M415 | LGE-WS-00397 | *H. m. malleti* | Florencia_Sucre | D-HW; D-FW; V-HW; V-FW |
| M418 | LGE-WS-00305 | *H. t. florencia* | Florencia_Sucre | D-HW; D-FW; V-HW; V-FW |
| M426 | LGE-WS-00361 | *H. m. malleti* | Florencia_Sucre | D-HW; D-FW; V-HW; V-FW |
| M433 | LGE-WS-00390 | *H. m. malleti* | Florencia_Sucre | V-HW; V-FW |
| M434 | LGE-WS-00363 | *H. m. malleti* | Florencia_Sucre | V-HW; V-FW |
| M451 | LGE-WS-00304 | *H. t. florencia* | Florencia_Sucre | D-HW; D-FW; V-HW; V-FW |
| M462 | LGE-WS-00337 | *H. t. florencia* | Florencia_Sucre | D-HW; D-FW; V-HW; V-FW |
| M468 | LGE-WS-00383 | *H. m. malleti* | Florencia_Sucre | D-HW; D-FW; V-HW; V-FW |
| M471 | LGE-WS-00307 | *H. t. florencia* | Florencia_Sucre | D-HW; D-FW; |
| M472 | LGE-WS-00311 | *H. t. florencia* | Florencia_Sucre | D-HW; D-FW; V-HW; V-FW |
| M474 | LGE-WS-00359 | *H. m. malleti* | Florencia_Sucre | V-HW; V-FW |
| M583 | LGE-WS-00367 | *H. m. malleti* | Florencia | D-HW; D-FW; V-HW; V-FW |
| M584 | LGE-WS-00357 | *H. m. malleti* | Florencia_Sucre | D-HW; D-FW |
| M587 | LGE-WS-00306 | *H. t. florencia* | Florencia_Sucre | D-HW; D-FW; V-HW; V-FW |
| M588 | LGE-WS-00310 | *H. t. florencia* | Florencia_Sucre | D-HW; D-FW; V-HW; V-FW |
| M589 | LGE-WS-00370 | *H. m. malleti* | Florencia_Sucre | D-HW; D-FW; V-HW; V-FW |
| M590 | LGE-WS-00395 | *H. m. malleti* | Florencia_Sucre | D-HW; D-FW; V-HW; V-FW |
| M592 | LGE-WS-00394 | *H. m. malleti* | Florencia_Sucre | D-HW; D-FW; V-HW; V-FW |
| M593 | LGE-WS-00309 | *H. t. florencia* | Florencia_Sucre | D-HW; D-FW; V-HW; V-FW |
| M594 | LGE-WS-00368 | *H. m. malleti* | Florencia | D-HW; D-FW; V-HW; V-FW |
| M595 | LGE-WS-00318 | *H. t. florencia* | Florencia_Sucre | D-HW; D-FW; V-HW; V-FW |
| M596 | LGE-WS-00308 | *H. t. florencia* | Florencia_Sucre | D-HW; D-FW; V-HW; V-FW |
| M598 | LGE-WS-00377 | *H. m. malleti* | Florencia_Sucre | D-HW; D-FW; V-HW; V-FW |
| M602 | LGE-WS-00303 | *H. t. florencia* | Florencia_Sucre | D-HW; D-FW; V-HW; V-FW |
| M606 | LGE-WS-00352 | *H. m. malleti* | Florencia_Sucre | D-HW; D-FW; V-HW; V-FW |
| M607 | LGE-WS-00317 | *H. t. florencia* | Florencia_Sucre | D-HW; D-FW; V-HW; V-FW |
| M610 | LGE-WS-00379 | *H. m. malleti* | Florencia_Sucre | D-HW; D-FW; V-HW; V-FW |
| M611 | LGE-WS-00316 | *H. t. florencia* | Florencia_Sucre | D-HW; D-FW; V-HW; V-FW |
| M612 | LGE-WS-00319 | *H. t. florencia* | Florencia_Sucre | D-HW; D-FW; V-HW; V-FW |
| M614 | LGE-WS-00355 | *H. m. malleti* | Florencia_Sucre | D-HW; D-FW; V-HW; V-FW |
| M616 | LGE-WS-00334 | *H. t. florencia* | Florencia_Sucre | D-HW; D-FW; V-HW; V-FW |
| M618 | LGE-WS-00324 | *H. t. florencia* | Florencia_Sucre | D-HW; D-FW |
| M620 | LGE-WS-00302 | *H. t. florencia* | Florencia_Sucre | D-HW; D-FW; V-HW; V-FW |
| M622 | LGE-WS-00332 | *H. t. florencia* | Florencia_Sucre | D-HW; D-FW |
| M1009 | LGE-WS-00333 | *H. t. florencia* | Florencia_Sucre | D-HW; D-FW; V-HW; V-FW |
| M1010 | LGE-WS-00313 | *H. t. florencia* | Florencia_Sucre | D-HW; D-FW; V-HW; V-FW |
| M1016 | LGE-WS-00378 | *H. m. malleti* | Florencia_Sucre | D-HW; D-FW; V-HW; V-FW |
| M1074 | LGE-WS-00315 | *H. t. florencia* | Florencia_Sucre | D-HW; D-FW; V-HW; V-FW |
| M1075 | LGE-WS-00314 | *H. t. florencia* | Florencia_Sucre | D-HW; D-FW; V-HW; V-FW |
| M1079 | LGE-WS-00348 | *H. t. florencia* | Florencia_Sucre | D-HW; D-FW; V-HW; V-FW |
| M1084 | LGE-WS-00328 | *H. t. florencia* | Florencia_Sucre | D-HW; D-FW; V-HW; V-FW |
| M1085 | LGE-WS-00329 | *H. t. florencia* | Florencia_Sucre | D-HW; D-FW |
| M1094 | LGE-WS-00330 | *H. t. florencia* | Florencia_Sucre | D-HW; D-FW; V-HW; V-FW |
| M1098 | LGE-WS-00396 | *H. m. malleti* | Florencia_Sucre | D-HW; D-FW; V-HW; V-FW |
| M1196 | LGE-WS-00354 | *H. m. malleti* | Florencia_Paraiso | D-HW; D-FW; V-HW; V-FW |
| M1283 | LGE-WS-00353 | *H. m. malleti* | Florencia_Paraiso | D-HW; D-FW; V-HW; V-FW |
| M1288 | LGE-WS-00364 | *H. m. malleti* | Florencia_Paraiso | V-HW; V-FW |
| M1321 | LGE-WS-00365 | *H. m. malleti* | Florencia_Paraiso | D-HW; D-FW; V-HW; V-FW |
| M1441 | LGE-WS-00381 | *H. m. malleti* | Florencia_Sucre | D-HW; D-FW; V-HW; V-FW |
| M1507 | LGE-WS-00389 | *H. m. malleti* | Florencia_Paraiso | V-HW; V-FW |
| M1511 | LGE-WS-00387 | *H. m. malleti* | Florencia_Paraiso | D-HW; D-FW; V-HW; V-FW |
| M1512 | LGE-WS-00386 | *H. m. malleti* | Florencia_Paraiso | D-HW; D-FW; V-HW; V-FW |
| M1514 | LGE-WS-00384 | *H. m. malleti* | Florencia_Paraiso | D-HW; D-FW; V-HW; V-FW |
| M1522 | LGE-WS-00385 | *H. m. malleti* | Florencia_Paraiso | D-HW; D-FW; V-HW; V-FW |
| M1754 | LGE-WS-00331 | *H. t. florencia* | Florencia_Sucre | D-HW; D-FW; V-HW; V-FW |
| M1757 | LGE-WS-00376 | *H. m. malleti* | Florencia_Sucre | D-HW; D-FW; V-HW; V-FW |
| M1758 | LGE-WS-00345 | *H. t. florencia* | Florencia_Sucre | D-HW; D-FW; V-HW; V-FW |
| M1767 | LGE-WS-00391 | *H. m. malleti* | Florencia_Sucre | D-HW; D-FW; V-HW; V-FW |
| M1769 | LGE-WS-00343 | *H. t. florencia* | Florencia_Sucre | D-HW; D-FW; V-HW; V-FW |
| M1770 | LGE-WS-00388 | *H. m. malleti* | Florencia_Sucre | V-HW; V-FW |
| M1771 | LGE-WS-00344 | *H. t. florencia* | Florencia_Sucre | D-HW; D-FW; V-HW; V-FW |
| M1772 | LGE-WS-00321 | *H. t. florencia* | Florencia_Sucre | D-HW; D-FW; V-HW; V-FW |
| M1773 | LGE-WS-00369 | *H. m. malleti* | Florencia_Sucre | D-HW; D-FW; V-HW; V-FW |
| M1774 | LGE-WS-00360 | *H. m. malleti* | Florencia_Sucre | D-HW; D-FW; V-HW; V-FW |
| M1805 | LGE-WS-00320 | *H. t. florencia* | Florencia_Sucre | D-HW; D-FW; V-HW; V-FW |
| M1808 | LGE-WS-00322 | *H. t. florencia* | Florencia_Sucre | D-HW; D-FW |
| M1813 | LGE-WS-00366 | *H. m. malleti* | Florencia | D-HW; D-FW; V-HW; V-FW |
| M1814 | LGE-WS-00399 | *H. m. malleti* | Florencia_Sucre | D-HW; D-FW |
| M1817 | LGE-WS-00323 | *H. t. florencia* | Florencia_Sucre | V-HW; V-FW |
| M1823 | LGE-WS-00356 | *H. m. malleti* | Florencia_Paraiso | D-HW; D-FW; V-HW; V-FW |
| M1845 | LGE-WS-00362 | *H. m. malleti* | Florencia_Paraiso | D-HW; D-FW; V-HW; V-FW |
| M1846 | LGE-WS-00336 | *H. t. florencia* | Florencia_Sucre | D-HW; D-FW; V-HW; V-FW |
| M2347 | LGE-WS-00393 | *H. m. malleti* | Florencia_Paraiso | D-HW; D-FW; V-HW; V-FW |
| M2360 | LGE-WS-00374 | *H. m. malleti* | Florencia_Paraiso | D-HW; D-FW; V-HW; V-FW |
| M2408 | LGE-WS-00382 | *H. m. malleti* | Florencia_Paraiso | D-HW; D-FW; V-HW; V-FW |
| M3544 | LGE-WS-00358 | *H. m. malleti* | Florencia_Paraiso | D-HW; D-FW; V-HW; V-FW |
| M3765 | LGE-WS-00347 | *H. t. florencia* | Florencia_Sucre | D-HW; D-FW; V-HW; V-FW |
| M3767 | LGE-WS-00339 | *H. t. florencia* | Florencia_Sucre | D-HW; D-FW; V-HW; V-FW |
| M3874 | LGE-WS-00341 | *H. t. florencia* | Florencia_Sucre | D-HW; D-FW; V-HW; V-FW |
| M3875 | LGE-WS-00327 | *H. t. florencia* | Florencia_Sucre | D-HW; D-FW |

Table S2. Female behavioural response towards males with normal and altered wing phenotype after testing the importance of wing colour pattern in mate preference. Behaviours are classified as Acceptance or Rejection. Asterisk (*) is indicative of statistically significance (α=0.01) according to GLMM.

| **Behaviour** | | ***H. m. malleti*** | ***H. t. florencia*** |
| --- | --- | --- | --- |
| Flutter | Acceptance | p=0.4213 | p=0.02956 |
| Fly towards | Acceptance | p=0.349 | p=0.7642 |
| Slow flat | Acceptance | p=0.4352 | p=0.2687 |
| Wings open | Acceptance | p=0.1576 | p=0.008592* |
| Abdomen exposed | Acceptance | p=0.6427 | p=0.7956 |
| Fly away | Rejection | p=0.125 | p=0.2246 |
| Tucked up | Rejection | p=0.2706 | p=0.0184 |
| Erratic flutter | Rejection | p=0.1892 | p=0.1142 |
| Abdomen bent | Rejection | p=0.7841 | p=0.01848 |

Table S3. Female behavioural responses in triads that tested female preference for males “perfumed” with a hexanic extract from five males either of *H. m. malleti* or *H. t. florencia*. Asterisk (*) is indicative of statistically significance (α=0.01) according to GLMM.

| **Behaviour** | | ***H. m. malleti*** | ***H. t. florencia*** |
| --- | --- | --- | --- |
| Flutter | Acceptance | p=8,72e-13* | p=1,42e-06* |
| Fly towards | Acceptance | p=2.20e-16* | p=7,79e-08* |
| Slow flat | Acceptance | p=2,97e-11* | p=2,79e-11* |
| Wings open | Acceptance | p=2.20e-16* | p=2.20e-16* |
| Abdomen exposed | Acceptance | p=2.20e-16* | p=7,27e-14* |
| Fly away | Rejection | p=2,87e-11* | p=2.20e-16* |
| Tucked up | Rejection | p=2,23e-04* | p=1,64e-02* |
| Erratic flutter | Rejection | p=4,16e-11* | p=2.20e-16* |
| Abdomen bent | Rejection | p=1,83e-07* | p=4,96e-11* |

Table S4. Amount (ng) of compounds remained in the wings of perfumed males with the heterospecific hexanic extract before evaporation. “pure species” indicate the mean of the compound present in the pure species blend. We dissected the wings after 1 minute, 30 minutes and 60 minutes after spreading the heterospecific hexanic extract. RI, retention index.

| **Name** | **RI** | ***H. melpomene malleti*** | | | | ***H. timareta florencia*** | | | |
| --- | --- | --- | --- | --- | --- | --- | --- | --- | --- |
|  |  | **Pure species** | **1 minute** | **30 minutes** | **60 minutes** | **Pure species** | **1 minute** | **30 minutes** | **60 minutes** |
| Unknown | 958.50 | 0.14 | 0.26 | 0.00 | 0.00 | - | - | - | - |
| Limonene | 1023.60 | - | - | - | - | 0.29 | 0.20 | 0.00 | 0.00 |
| Phenylacetaldehyde | 1036.10 | - | - | - | - | 0.17 | 0.00 | 0.00 | 0.00 |
| Methyl salicylate | 1187.20 | - | - | - | - | 1.44 | 1.37 | 0.00 | 0.03 |
| Dodecane | 1117.30 | - | - | - | - | 2.78 | 2.82 | 2.01 | 1.15 |
| Unknown | 1174.80 | 0.20 | 0.14 | 0.14 | 0.41 | 0.45 | 2.35 | 2.20 | 0.83 |
| (Z)-3-Hexenyl isobutyrate | 1233.30 | 19.22 | 17.46 | 0.00 | 0.00 | - | - | - | - |
| Hexyl 3-methylbutyrate | 1239.50 | 53.33 | 0.00 | 0.00 | 0.00 | - | - | - | - |
| Unknown | 1243.70 | 0.58 | 0.30 | 0.00 | 0.00 | - | - | - | - |
| Alkane | 1265.40 | - | - | - | - | 1.82 | 0.46 | 0.00 | 0.00 |
| Tridecane | 1300.00 | 15.16 | 18.51 | 15.01 | 5.62 | 7.67 | 7.24 | 4.64 | 1.72 |
| Tetradecane | 1302.40 | - | - | - | - | 0.67 | 0.70 | 0.64 | 0.62 |
| 5-Decanolide | 1369.50 | - | - | - | - | 4.80 | 4.76 | 4.44 | 1.16 |
| alpha-Copaene | 1371.50 | 1.31 | 0.00 | 0.00 | 0.00 | - | - | - | - |
| Dihydroactinidiolide | 1391.70 | 17.67 | 17.35 | 14.13 | 8.36 | 14.40 | 13.25 | 7.91 | 6.63 |
| Unknown | 1394.80 | 0.90 | 0.73 | 0.41 | 0.00 | - | - | - | - |
| Ethyl 4-ethoxybenzoate | 1402.60 | 8.29 | 9.40 | 1.03 | 0.56 | 12.09 | 15.41 | 14.39 | 14.06 |
| Homovanillyl alcohol | 1412.10 | 1.56 | 1.68 | 1.16 | 0.45 | 0.25 | 0.00 | 0.00 | 0.00 |
| Methy 4-hydroxybenzoate | 1449.00 | 2.51 | 2.10 | 0.00 | 0.00 | - | - | - | - |
| Methyl 3,4-dimethoxybenzoate | 1464.10 | 8.97 | 7.53 | 5.88 | 3.25 | - | - | - | - |
| Unknown | 1470.10 | 0.92 | 0.57 | 0.56 | 0.30 | - | - | - | - |
| Unknown | 1488.30 | - | - | - | - | 0.11 | 0.00 | 0.00 | 0.00 |
| Unknown | 1495.20 | 0.10 | 0.16 | 0.12 | 0.06 | - | - | - | - |
| Syringaaldehyde | 1519.00 | 291.80 | 293.20 | 280.37 | 251.10 | 277.15 | 263.46 | 207.00 | 174.30 |
| 3,5-Dimethoxy 4-hydroxybenzyl alcohol | 1565.00 | - | - | - | - | 7.17 | 0.00 | 0.00 | 0.00 |
| Propyl 4-hydroxybenzoate | 1614.60 | - | - | - | - | 2.70 | 0.00 | 0.00 | 0.00 |
| Methyl 1H-indol-3-carboxylate | 1663.10 | - | - | - | - | 0.24 | 0.28 | 0.26 | 0.07 |
| Unknown | 1715.80 | 1.22 | 2.10 | 0.00 | 0.64 | - | - | - | - |
| Tricosene | 1740.20 | - | - | - | - | 1.13 | 1.27 | 1.23 | 0.56 |
| Hexadecanoic acid | 1818.00 | - | - | - | - | 1.75 | 1.46 | 0.61 | 0.31 |
| Octadecanal | 1868.00 | 1094.90 | 826.74 | 202.60 | 130.10 | - | - | - | - |
| Isopropyl Palmitate | 1877.50 | - | - | - | - | 1.93 | 2.62 | 2.04 | 1.60 |
| Unknown | 1922.10 | 16.56 | 22.88 | 21.86 | 0.80 | - | - | - | - |
| Heneicosene | 1946.10 | - | - | - | - | 12.30 | 3.45 | 0.00 | 0.00 |
| Heneicosane | 1952.10 | 298.84 | 325.10 | 287.02 | 171.70 | 174.57 | 193.60 | 165.48 | 146.40 |
| 1-Octadecanol | 1929.60 | 27.83 | 34.65 | 34.44 | 21.10 | 26.23 | 15.28 | 9.59 | 0.00 |
| (Z)-11-Eicosenal | 2031.20 | 231.99 | 63.82 | 59.68 | 32.72 | 3.86 | 2.48 | 0.00 | 0.00 |
| Docosane | 2044.40 | 0.28 | 4.70 | 4.41 | 0.00 | - | - | - | - |
| Eicosanal | 2058.90 | 26.43 | 35.88 | 31.85 | 0.00 | - | - | - | - |
| Unknown | 2109.90 | - | - | - | - | 0.50 | 0.00 | 0.00 | 0.00 |
| Unknown | 2135.80 | 1.54 | 1.99 | 1.77 | 0.64 | 0.20 | 0.25 | 0.23 | 0.11 |
| Tricosane | 2137.20 | 27.27 | 0.00 | 0.00 | 0.00 | 3.36 | 0.00 | 0.00 | 0.00 |
| Isopropyl oleate | 2187.70 | - | - | - | - | 0.00 | 0.03 | 0.00 | 0.00 |
| Ethyl stearate | 2190.40 | - | - | - | - | 1.77 | 1.24 | 0.00 | 0.00 |
| (Z)-13-Docosenal | 2217.80 | 23.67 | 0.00 | 0.00 | 0.00 | - | - | - | - |
| Eicosane | 2228.10 | 2.65 | 1.34 | 2.17 | 5.26 | 2.56 | 3.27 | 2.36 | 1.83 |
| Heptacosane | 2489.80 | 60.12 | 78.75 | 79.10 | 219.00 | 31.49 | 30.28 | 21.81 | 11.74 |
| Octacosane | 2569.30 | - | - | - | - | 0.40 | 0.00 | 0.00 | 0.00 |
| Hexacosanal | 2594.80 | - | - | - | - | 0.66 | 0.00 | 0.00 | 0.00 |
| Nonacosane | 2652.30 | 25.34 | 26.21 | 24.79 | 0.00 | 14.03 | 11.58 | 2.88 | 1.34 |
| 13,17-Dimethylnonacosane | 2688.10 | - | - | - | - | 1.83 | 0.89 | 0.00 | 0.00 |
| Ethyl benzoate | 2686.30 | - | - | - | - | 0.31 | 0.32 | 0.30 | 0.00 |
| Octacosanal | 2740.80 | 12.51 | 15.64 | 13.83 | 0.00 | 28.55 | 24.31 | 11.60 | 1.18 |
| Cholesterol | 2807.00 | 286.19 | 229.63 | 128.30 | 0.00 | 236.60 | 92.43 | 87.18 | 38.70 |
| Hentriacontane | 2807.80 | 58.10 | 53.88 | 47.26 | 0.00 | 115.60 | 65.82 | 18.58 | 15.55 |
| 13,17-Dimethylhentriacontane | 2835.80 | 12.95 | 10.76 | 1.64 | 0.00 | 3.93 | 2.65 | 1.74 | 0.00 |
| 2-Eicosyl-5-nonyltetrahydrofuran | 3121.50 | - | - | - | - | 3.89 | 0.00 | 0.00 | 0.00 |

Table S5. Behavioural response in F_1_ and BC females towards pure males of *H. m. malleti* and *H. t. florencia*. Asterisk (*) is indicative of statistically significance (α=0.01) according to GLMM.

| **Behaviour** | | **F_1_ Females** | **BC Females** |
| --- | --- | --- | --- |
| Flutter | Acceptance | p=0,147 | p=0,75 |
| Fly towards | Acceptance | p=0,0743 | p=9,90e-10* |
| Slow flat | Acceptance | p=9,64e-13* | p=2,20e-16* |
| Wings open | Acceptance | p=3,30e-15* | p=3,91e-14* |
| Abdomen exposed | Acceptance | p=1,10e-03* | p=2,20e-16* |
| Fly away | Rejection | p=0,439 | p=2,20e-16* |
| Tucked up | Rejection | p=2,57e-09* | p=2,20e-16* |
| Erratic flutter | Rejection | p=3,11e-11* | p=2,20e-16* |
| Abdomen bent | Rejection | p=6,91e-07* | p=0,03141 |

Table S6. Compounds identified in extracts of wing androconia of *H. melpomene malleti*, *H. timareta florencia*, F_1_ and backcrosses males. RI, retention index. Mean ± SD amounts in ng.

| **Name** | **RI** | ***H. melpomene malleti*** | ***H. timareta florencia*** | **F1** | **BC** |
| --- | --- | --- | --- | --- | --- |
|  |  | **Mean ± SD** | **Mean ± SD** | **Mean ± SD** | **Mean± SD** |
| Unknown | 902.60 | - | - | 3.80 ± 4.91 | 0.78 ± 1.60 |
| Dimethyl sulfone | 916.00 | - | - | 0.76 ± 1.33 | 0.50 ± 1.99 |
| Unknown | 958.50 | 0.13 ± 0.76 | 0.43 ± 1.69 | 7.15 ± 4.04 | 11.25 ± 5.68 |
| Limonene | 1023.60 | - | 0.28 ± 0.52 | - | - |
| Phenylacetaldehyde | 1036.10 | - | 0.16 ± 0.77 | - | - |
| Unknown | 1037.60 | - | - | 1.50 ± 2.40 | 1.16 ± 1.85 |
| Nonanal | 1100.70 | 0.58 ± 1.51 | 1.47 ± 2.94 | 3.67 ± 4.97 | - |
| Dodecane | 1117.30 | - | 2.91 ± 5.72 | - | - |
| Unknown | 1174.80 | - | 0.06 ± 3.09 | - | - |
| Methyl salicylate | 1187.20 | 2.44 ± 2.51 | 1.39 ± 1.99 | - | - |
| Decanal | 1200.90 | - | - | 3.20 ± 8.32 | - |
| (Z)-3-Hexenyl isobutyrate | 1233.30 | 19.21 ± 14.04 | - | - | - |
| Hexyl 3-methylbutyrate | 1239.50 | 53.33 ± 144.25 | - | - | - |
| Alkane | 1265.40 | - | 1.76 ± 9.72 | - | - |
| Tridecane | 1300.00 | 15.16 ± 12.07 | 8.15 ± 9.15 | - | - |
| Tetradecane | 1302.90 | - | 0.92 ± 2.54 | - | - |
| 5-Decanolide | 1369.90 | - | 4.65 ± 5.40 | 1.06 ± 1.88 | 0.12 ± 0.64 |
| alpha-Copaene | 1371.50 | 1.31 ± 1.88 | - | - | - |
| Dihydroactinidiolide | 1392.90 | 17.67 ± 11.54 | 13.96 ± 15.69 | 1.71 ± 1.84 | 0.93 ± 2.02 |
| Ethyl 4-ethoxybenzoate | 1403.40 | 8.28 ± 6.87 | 12.12 ± 16.23 | 8.28 ± 5.88 | 5.65 ± 6.65 |
| Homovanillin alcohol | 1413.10 | 1.55 ± 2.93 | 0.249 ± 0.78 | 0.27 ± 0.53 | 0.12 ± 0.47 |
| 6,10-Dimethyl 5,9-undecadien-2-one | 1447.60 | 2.64 ± 3.10 | - | - | - |
| Methy 4-hydroxybenzoate | 1449.00 | 2.51 ± 4.83 | - | - | - |
| Methyl 3,4-dimethoxybenzoate | 1465.90 | 8.97 ± 18.25 | - | - | - |
| Unknown | 1470.10 | 0.19 ± 0.50 | - | - | 0.33 ± 0.65 |
| Unknown | 1488.30 | - | 0.09 ± 0.46 | - | - |
| Unknown | 1495.20 | - | - | 0.06 ± 0.22 | - |
| Syringaaldehyde | 1519.00 | 291.8 ± 234.84 | 268.75 ± 285.36 | 56.11 ± 62.01 | 34.18 ± 50.06 |
| 3,5-Dimethoxy 4-hydroxybenzyl alcohol | 1565.00 | - | 6.95 ± 36.05 | - | - |
| Propyl 4-hydroxybenzoate | 1614.60 | - | 3.02 ± 5.43 | - | - |
| Methyl 1H-indol-3-acetate | 1663.10 | - | 0.23 ± 0.75 | - | - |
| Unknown | 1715.80 | 0.89 ± 1.51 | - | 2.60 ± 3.14 | 1.57 ± 1.77 |
| Ethyl benzoate | 1765.30 | - | 0.30 ± 0.89 | 0.19 ± 0.53 | - |
| 16-Hexadecanolide | 1769.10 | - | - | - | 0.92 ± 4.01 |
| Unknown | 1795.00 | - | 3.72 ± 20.66 | 0.16 ± 0.40 | - |
| Hexadecanoic acid | 1826.50 | - | 2.76 ± 8.24 | - | - |
| Octadecanal | 1869.80 | 1094.91 ± 519.64 | 1.40 ± 5.84 | 19.25 ± 35.04 | 1.86 ± 5.38 |
| Unknown | 1873.80 | - | - | 0.14 ± 0.48 | 1.64 ± 4.85 |
| Isopropyl palmitate | 1878.30 | - | 1.86 ± 6.02 | - | 3.00 ± 8.28 |
| Unknown | 1923.40 | 16.56 ± 14.99 | - | - | - |
| Unknown | 1929.30 | 0.57 ± 2.42 | - | 1.02 ± 1.39 | - |
| 1-Octadecanol | 1929.60 | 27.83 ± 33.07 | 29.85 ± 138.42 | 2.82 ± 6.48 | - |
| Heneicosene | 1946.10 | 0.47 ± 1.90 | 26.80 ± 104.03 | 5.97 ± 18.76 | 4.22 ± 11.20 |
| Heneicosane | 1953.70 | 298.83 ± 335.05 | 174.57 ± 225.37 | 132.51 ± 102.24 | 125.07 ± 122.00 |
| (Z)-11-Eicosenal | 2032.70 | 231.98 ± 255.99 | 4.39 ± 11.48 | 22.25 ± 20.38 | 3.75 ± 9.35 |
| Docosane | 2045.50 | 0.27 ± 0.99 | 6.25 ± 22.60 | - | - |
| Eicosanal | 2058.90 | 43.43 ± 53.98 | - | 1.46 ± 3.12 | - |
| Unknown | 2088.50 | - | 1.93 ± 11.14 | - | - |
| Tricosene | 2106.80 | - | 2.84 ± 16.35 | 0.20 ± 0.68 | - |
| Unknown | 2109.90 | 0.91 ± 4.15 | - | 0.28 ± 0.72 | 0.37 ± 0.86 |
| Unknown | 2135.80 | 1.53 ± 4.23 | - | - | - |
| Tricosane | 2137.20 | 27.27 ± 151.83 | 8.46 ± 31.72 | - | - |
| Ethyl stearate | 2190.40 | - | 1.71 ± 3.18 | - | 0.05 ± 0.19 |
| (Z)-13-Docosenal | 2217.80 | 23.66 ± 68.98 | - | 0.62 ± 2.08 | - |
| Docosane | 2227.90 | - | 0.19 ± 0.92 | 1.75 ± 3.75 | 0.36 ± 0.86 |
| Eicosane | 2228.10 | 2.63 ± 5.25 | 3.43 ± 7.06 | 0.10 ± 0.35 | 0.23 ± 0.56 |
| Unknown | 2321.80 | 0.10 ± 0.57 | - | - | 0.98 ± 2.78 |
| Heptacosane | 2411.00 | 56.11 ± 73.64 | 12.16 ± 39.18 | 23.78 ± 17.25 | 36.12 ± 30.35 |
| Octacosane | 2569.30 | - | 2.63 ± 15.11 | - | - |
| Hexacosanal | 2594.80 | - | 5.20 ± 26.32 | - | - |
| Methylheptacosane | 2596.10 | 15.88 ± 26.13 | 12.42 ± 39.54 | 4.24 ± 6.31 | 6.34 ± 15.06 |
| Nonacosane | 2652.30 | 25.34 ± 41.2 | 13.60 ± 25.45 | 31.43 ± 23.85 | 31.24 ± 38.05 |
| Unknown | 2687.70 | - | 0.39 ± 2.25 | 0.43 ± 1.44 | - |
| 13,17 Dimethylnonacosane | 2688.10 | - | 1.77 ± 4.84 | 7.26 ± 7.19 | 1.17 ± 3.04 |
| Octacosanal | 2740.80 | 12.50 ± 21.93 | 28.82 ± 52.43 | 28.71 ± 28.59 | 23.6 ± 24.67 |
| Unknown | 2794.20 | 1.21 ± 3.23 | - | 0.15 ± 0.52 | 21.78 ± 69.18 |
| Cholesterol | 2807.00 | 286.19 ± 445.59 | 250.36 ± 219.16 | 218.74 ± 95.74 | 204.56 ± 135.52 |
| Hentriacontane | 2807.80 | 58.09 ± 100.82 | 118.43 ± 206.82 | 88.60 ± 39.46 | 52.32 ± 32.19 |
| 13,17-Dimethylhentriacontane | 2835.80 | 12.95 ± 28.21 | 3.81 ± 17.50 | 18.13 ± 17.65 | 4.16 ± 8.97 |
| 2-Eicosyl-5-heptyltetrahydrofuran | 2923.70 | 97.19 ± 257.04 | - | - | - |
| 2-Eicosyl-5-nonyltetrahydrofuran | 3121.50 | 10.11 ± 56.30 | 4.69 ± 13.95 | - | 1.44 ± 7.23 |

Table S7. Compounds identified in extracts of abdominal glands of *H. melpomene malleti*, *H. timareta florencia*, F_1_ and backcrosses males. RI, retention index. Mean ± SD amounts in ng.

| **Name** | **RI** | ***H. melpomene malleti*** | ***H. timareta florencia*** | ***F1*** | ***BC*** |
| --- | --- | --- | --- | --- | --- |
|  |  | **Mean ± SD** | **Mean ± SD** | **Mean ± SD** | **Mean ± SD** |
| Unknown | 903.90 | - | 1.58 ± 3.99 | 7.08 ± 5.85 | 13.70 ± 12.24 |
| Dimethyl sulfone | 919.00 | - | 0.34 ± 1.22 | - | - |
| (Z)-beta-Ocimene | 1037.50 | 11899.84 ± 7633.07 | - | 142.08 ± 270.49 | 78.92 ± 243.28 |
| Phenylacetonitril_Benzylcyanid | 1039.80 | - | 62.59 ± 95.31 | - | - |
| (E)-beta-Ocimene | 1048.20 | 12096 ± 7193.57 | - | 2247.58 ± 3047.13 | 34.18 ± 75.04 |
| 2-sec-Butyl-3-methoxypyrazine | 1170.10 | - | 74.80 ± 91.67 | 12.45 ± 16.86 | 55.28 ± 34.86 |
| 2-isobutyl-3-methoxy pyrazine | 1177.70 | 0.02 ± 0.11 | - | 1.26 ± 1.51 | 2.43 ± 3.94 |
| Methyl salicylate | 1188.50 | - | 0.12 ± 0.54 | 1.74 ± 1.28 | 1.27 ± 1.68 |
| 5-Decanolide | 1203.00 | 0.42 ± 2.24 | 0.18 ± 1.03 | - | - |
| Dihydroedulan II | 1284.90 | 43.92 ± 40.09 | 5.52 ± 14.98 | 1.82 ± 2.26 | - |
| Tridecane | 1300.00 | 1.33 ± 1.91 | 0.44 ± 0.87 | - | - |
| Nonadecane | 1302.70 | 554.93 ± 1403.34 | 721.14 ± 3082.59 | - | - |
| alpha-Copaene | 1374.10 | 0.02 ± 0.14 | 2.53 ± 6.81 | - | - |
| GC-EAD active compound | 1395.20 | 264.14 ± 333.29 | 622.68 ± 661.95 | 122.10 ± 124.78 | 47.67 ± 66.71 |
| Dihydroactinidiolide | 1517.30 | 0.09 ± 0.48 | 21.34 ± 120.35 | - | - |
| Ethyl 4-ethoxybenzoate | 1520.20 | 8.81 ± 7.73 | 18.26 ± 40.40 | 9.51 ± 6.44 | 6.02 ± 7.69 |
| Homovanillyl alcohol | 1534.00 | 1.43 ± 2.86 | 1.40 ± 4.25 | 0.18 ± 0.48 | 1.38 ± 2.72 |
| Unknown | 1542.90 | 0.18 ± 1.00 | 0.49 ± 2.54 | 3.83 ± 5.72 | 0.55 ± 1.05 |
| Heptadecane | 1700.40 | 155.78 ± 602.03 | 0.13 ± 0.76 | 0.24 ± 0.82 | - |
| Benzyl_salicylate | 1708.20 | 0.34 ± 0.84 | - | - | - |
| Unknown | 1712.00 | - | 23.37 ± 69.08 | 0.76 ± 1.33 | - |
| 14-Tetradecanolide | 1721.50 | 5.86 ± 14.98 | 30.18 ± 62.64 | 40.03 ± 48.57 | 9.77 ± 15.12 |
| Hexadecatrienolide | 1726.10 | - | 3.14 ± 6.25 | - | - |
| 9_11-Hexadecadien-11-olide | 1734.70 | - | 13.31 ± 25.54 | - | - |
| Ethyl benzoate | 1764.60 | 6.86 ± 14.56 | 11.54 ± 20.45 | - | - |
| Unknown | 1771.70 | - | 1.06 ± 3.42 | 32.52 ± 50.73 | - |
| Macrolide | 1773.70 | 0.5423 ± 2.86 | 0.81 ± 2.44 | - | 0.34 ± 1.78 |
| (Z2_Z4)-C16-15-olide | 1806.60 | - | 1.75 ± 6.20 | - | - |
| Hexadecen-11-olide | 1819.30 | - | 12.32 ± 16.05 | - | - |
| Octadecatrienolide | 1823.40 | 32.10 ± 32.03 | 26.36 ± 36.51 | - | - |
| Hexadecenolide | 1857.00 | 11.18 ± 24.31 | 126.3 ± 214.21 | 6.04 ± 11.35 | 13.09 ± 23.92 |
| Macrolide | 1923.60 | 6.98 ± 36.97 | 2.85 ± 10.88 | - | 0.45 ± 2.33 |
| Heptadecanal | 1923.70 | - | - | 8.64 ± 12.30 | 12.93 ± 22.66 |
| 16-Hexadecanolide | 1924.80 | - | 124.38 ± 160.04 | 4.16 ± 11.61 | - |
| Brassicalactone | 1960.90 | - | 278.62 ± 496.97 | 11.79 ± 22.16 | 20.40 ± 50.00 |
| Octadecen-11-olide | 2002.70 | 2.26 ± 9.61 | 242.74 ± 436.73 | 5.09 ± 7.11 | 11.28 ± 22.48 |
| Eicosane | 2005.10 | 1.04 ± 3.16 | 1354.90 ± 1833.53 | 12.15 ± 27.11 | 10.83 ± 11.16 |
| Isopropyl palmitate | 2029.80 | 2.17 ± 9.03 | 75.63 ± 183.90 | 8.99 ± 13.25 | 11.63 ± 32.00 |
| (Z)-9-C18-11-olide | 2032.80 | 4.33 ± 10.11 | 841.29 ± 1138.35 | 319.30 ± 450.38 | 985.96 ± 1393.24 |
| (Z)-9-C18-13-olide | 2038.70 | 221.76 ± 278.84 | 1933.83 ± 2671.28 | 544.38 ± 452.76 | 1779.89 ± 1709.61 |
| 12-Octadecanolide | 2051.80 | - | 3.13 ± 7.20 | 0.54 ± 1.29 | 0.61 ± 2.44 |
| Macrolide | 2056.70 | 0.8512 ± 2.40 | 0.19 ± 1.12 | - | - |
| (E)-Octadec-9-en-12-olide | 2057.20 | - | 64.11 ± 79.56 | - | 25.10 ± 123.85 |
| Macrolide | 2058.90 | 5.31 ± 10.70 | 11.75 ± 22.80 | 3.83 ± 5.72 | 0.55 ± 1.05 |
| Isopropyl_octadecanoate | 2063.40 | 5.57 ± 16.87 | 277.67 ± 399.88 | - | - |
| (Z9,E11)-C18-13-olide | 2069.60 | 0.06 ± 0.35 | 1417.08 ± 3035.70 | 2.66 ± 7.33 | 165.54 ± 258.28 |
| Octadeca-9-11-dien-13-olide and 11-Octadecanolide | 2070.20 | - | 29.81 ± 101.85 | 36.17 ± 31.27 | 47.18 ± 166.82 |
| Isopropyl_linoleate | 2073.40 | 2.53 ± 7.92 | 929.63 ± 1218.84 | - | - |
| Henicosene | 2074.20 | 28.17 ± 40.01 | 20.59 ± 25.93 | 16.14 ± 15.61 | 327.90 ± 762.56 |
| 1-Octadecanol | 2081.50 | 116.92 ± 160.41 | 67.52 ± 138.34 | 16.10 ± 32.72 | - |
| Heneicosane | 2101.60 | 2122.79 ± 1879.48 | 1176.11 ± 2609.26 | 531.93 ± 392.53 | 610.33 ± 960.01 |
| Octadecen-18-olide | 2123.30 | 0.27 ± 1.47 | 15.88 ± 45.96 | 18.74 ± 29.46 | 20.29 ± 30.55 |
| 17-Octadecanolide | 2136.30 | - | 25.34 ± 143.36 | 109.12 ± 263.67 | 149.48 ± 501.09 |
| Isopropyl_octadecadienolate | 2130.20 | 1.67 ± 6.25 | 99.43 ± 369.98 | - | - |
| 9-Octadecen-18-olide | 2138.10 | 44.04 ± 122.04 | 159.13 ± 217.98 | 152.97 ± 190.59 | 190.42 ± 325.61 |
| Octadecanolide | 2158.50 | - | - | 6.56 ± 2.35 | - |
| Ethyl oleate | 2165.90 | 377.55 ± 474.37 | 606.93 ± 733.31 | 101.26 ± 172.42 | 0.34 ± 1.78 |
| Octadecadienolide | 2171.70 | 29.60 ± 71.78 | 700.86 ± 919.28 | 598.20 ± 448.49 | 1413.63 ± 1228.55 |
| Butyl hexadecanoate | 2186.90 | 9.05 ± 13.25 | 122.77 ± 197.54 | 28.75 ± 61.82 | 46.27 ± 79.45 |
| Isopentyl octadecadienoate | 2189.30 | - | 402.28 ± 1692.33 | 32.41 ± 48.63 | 25.33 ± 53.00 |
| Isopropyl oleate | 2196.10 | 969.26 ± 828.53 | 7715.08 ± 6651.53 | 2070.673 ± 1733.78 | 3285.38 ± 4207.11 |
| Docosane | 2200.90 | 3.00 ± 7.09 | 39.28 ± 124.92 | 6.32 ± 13.64 | 1.74 ± 4.54 |
| Butyl_octadecanoate | 2209.00 | 17.85 ± 27.85 | 115.71 ± 162.59 | - | - |
| Eicosanal | 2222.30 | 0.42 ± 2.24 | 1.05 ± 5.98 | 0.48 ± 1.60 | 0.70 ± 3.36 |
| Unknown | 2248.70 | 8.56 ± 20.40 | 19.21 ± 26.10 | 0.39 ± 1.29 | 0.23 ± 1.18 |
| 13-Eicosanolide | 2252.30 | - | 6.53 ± 18.44 | 33.26 ± 62.74 | 40.95 ± 71.75 |
| Tricosene | 2274.90 | 23.00 ± 33.65 | 45.26 ± 73.63 | 19.37 ± 23.37 | 30.76 ± 45.55 |
| Isobutyl oleate | 2297.70 | 243.89 ± 916.10 | 883.52 ± 1294.80 | 146.69 ± 205.08 | 51.79 ± 163.49 |
| Tricosane | 2303.20 | 168.89 ± 166.76 | 191.35 ± 273.22 | 61.88 ± 84.02 | 59.88 ± 67.54 |
| 2-Heneicosanol | 2310.60 | - | 196.39 ± 488.74 | 7.84 ± 26.02 | - |
| 11-Icosenol | 2317.40 | 11.18 ± 19.08 | 1260.80 ± 4099.99 | 2.66 ± 6.30 | 2.09 ± 5.97 |
| Butyl oleate | 2359.30 | 539.28 ± 615.35 | 2633.57 ± 3949.56 | 1805.67 ± 1680.41 | 3411.09 ± 4616.37 |
| Hexenyl hexadecanoate | 2379.10 | - | 12.53 ± 29.20 | 4.44 ± 10.55 | 10.27 ± 14.82 |
| Tetracosane | 2405.50 | 15.69 ± 68.45 | 104.19 ± 513.61 | 1.60 ± 5.31 | - |
| 1_3-Docosanediol | 2409.10 | 0.08 ± 0.32 | 17.25 ± 42.61 | - | - |
| Macrolide | 2417.30 | 2.324 ± 8.02 | 2.17 ± 11.45 | 0.76 ± 1.33 | - |
| Macrolide | 2418.70 | 2.318 ± 3.72 | 11.22 ± 16.59 | 19.73 ± 37.04 | 63.50 ± 97.01 |
| Isoprenyl octadec-11-enoate | 2433.60 | - | 92.19 ± 313.63 | - | - |
| Unknown | 2434.60 | 5.90 ±17.08 | 17.35 ± 51.73 | 0.39 ± 1.29 | 0.23 ± 1.18 |
| (Z)-13-Docosen-1-ol | 2461.40 | 87.02 ± 176.14 | 297.51 ± 425.23 | 39.84 ± 63.69 | 26.27 ± 71.84 |
| Eicosenolide | 2475.20 | - | - | 26.80 ± 44.29 | 58.17 ± 102.54 |
| 1- Docosanol | 2488.10 | 37.55 ± 57.04 | 140.36 ± 331.25 | 108.25 ± 126.81 | 137.70 ± 162.27 |
| Pentacosane | 2500.80 | 136.3 ± 141.58 | 206.84 ± 803.27 | 33.49 ± 39.19 | 30.39 ± 37.85 |
| (Z)-9-Tricosene | 2514.30 | - | 17.94 ± 38.29 | - | - |
| 11-methylpentacosane | 2532.50 | 48.70 ± 52.49 | 7.45 ± 22.8 | - | - |
| Docosen-22-olide | 2537.50 | - | 92.32 ± 117.00 | 38.40 ± 69.37 | - |
| Hexyl octadecadienoate | 2538.70 | - | 11.25 ± 35.99 | 7.01 ± 23.26 | 22.09 ± 37.33 |
| Hexyl octadecenoate and Hexenyl octadecenoate | 2553.20 | 114.56 ± 243.75 | 679.70 ± 974.09 | 586.15 ± 580.31 | 1173.90 ± 1111.75 |
| Hexenyl octadecatrienoate and Hexenyl octadecatrienoate | 2555.40 | - | 16.35 ± 41.97 | 41.83 ± 96.64 | 157.85 ± 196.27 |
| Benzyl hexadecanoate | 2571.20 | - | 12.51 ± 26.71 | 1.20 ± 4.00 | 1.10 ± 4.26 |
| Hexenyl octadecanoate | 2580.30 | 6.08 ± 16.75 | 4.99 ± 9.47 | 1.53 ± 2.67 | 2.33 ± 5.92 |
| Hexyl octadecanoate | 2594.70 | 0.743 ± 3.93 | 81.33 ± 95.56 | 25.69 ± 38.66 | 68.48 ± 95.79 |
| Hexacosane | 2601.10 | 5.54 ± 11.89 | 179.42 ± 1012.18 | 6.31 ± 15.96 | - |
| Tetracosenol | 2666.20 | 205.96 ± 239.41 | 220.54 ± 319.25 | 439.32 ± 530.86 | 405.506 ± 481.61 |
| 1-Tetracosanol | 2691.80 | 9.04 ± 21.35 | 47.67 ± 116.63 | 138.60 ± 309.57 | 45.47 ± 73.74 |
| Heptacosane | 2700.90 | 19.83 ± 47.97 | 225.04 ± 1162.37 | 98.30 ± 85.46 | 129.81 ± 143.60 |
| Tetracosenolide | 2735.80 | 2.96 ± 7.36 | 42.87 ± 56.81 | 33.63 ± 55.08 | 59.24 ± 128.91 |
| 1,3-Tetracosanediol | 2811.90 | - | - | 58.20 ± 120.24 | 8.24 ± 19.93 |
| Unknown | 2869.20 | 8.19 ± 19.09 | 0.29 ± 1.51 | 3.56 ± 9.01 | 3.15 ± 16.12 |
| Hexacosanal | 2871.80 | 8.43 ± 28.03 | 28.60 ± 49.18 | - | - |
| Nonacosane | 2901.50 | 1.57 ± 6.43 | 167.65 ± 924.25 | 15.15 ± 19.06 | 20.82 ± 35.97 |
| 13,17-Dimethylnonacosane | 2959.20 | - | 32.82 ± 147.47 | 43.77 ± 115.55 | 6.94± 20.82 |
| Cholesterol | 3099.40 | 192.05 ± 273.18 | 153.25 ± 263.26 | 617.34 ± 599.42 | 700.25 ± 647.53 |
| Unknown | 3147.40 | 0.08 ± 0.33 | 0.68 ± 3.85 | 6.26 ± 16.19 | - |
| 13,17-Dimethylhentriacontane | 3158.20 | 153.09 ± 173.73 | 395.95 ± 485.06 | 438.67 ± 328.29 | 349.94 ± 422.70 |
| 2-Nonyl-5-octadecyltetrahydrofuran | 3177.50 | 0.08 ± 0.33 | 0.20 ± 0.82 | 22.65 ± 57.78 | - |
| Campesterol or Ergostenol | 3207.30 | 1.42 ± 5.61 | 7.37 ± 17.36 | 16.35 ± 21.95 | 56.38 ± 120.37 |
| 13,17-Dimethyltritriacontane | 3348.00 | 24.37 ± 39.54 | 42.23 ± 105.73 | 93.10 ± 75.05 | 64.86 ± 88.28 |
| 2-Eicosyl-5-nonyltetrahydrofuran | 3370.90 | 14.77 ± 35.28 | 20.68 ± 48.38 | 63.61 ± 50.82 | 39.84 ± 72.41 |

### Table S8. Probability of mating in no-choice experiments.

Hmm: *H. m. malleti*; Htf: *H. t. florencia*; F1: Htf x Hmm; BC: backcrosses [Htf x Hmm] x Htf. Cross type is specified as female x male. [Confidence interval at 95%]. No-choice mating data was collected as was previously done in other Heliconius studies (1,2). This information allowed us to complete what is known about species premating barriers (3). Mating probability for interspecific and hybrid trials was obtained by maximizing the loge of the likelihood function (for details see 4,5).

| **Cross type** | | **N trials** | **Mating probability** | **Confidence interval** | **Source** |
| --- | --- | --- | --- | --- | --- |
| Control (conspecific) | Htf x Htf | 45 | 0.911 | [0.82 - 0.971] | Merot 2017 |
|  | Hmm x Hmm | 35 | 0.857 | [0.737 - 0.946] | Merot 2017 |
| Interspecific | Hmm x Htf | 13 | 0.152 | [0.04 - 0.363] | This study |
|  | Htf x Hmm | 16 | 0.188 | [0.157 - 0.377] | This study |
| Hybrid crosses | **F_1_ x Hmm** | **18** | **0** | **[0 - 0.024]** | **Merot 2017** |
|  | **F_1_ x Htf** | **24** | **0.249** | **[0.119 - 0.4]** | **Merot 2017** |
|  | Hmm x F_1_ | 8 | 0 | [0 - 0.011] | Merot 2017 |
|  | Htf x F_1_ | 10 | 0.2 | [0.04 - 0.45] | Merot 2017 |
|  | F_1_ x F_1_ | 4 | 0 | [0 - 0.0055] | This study |
|  | **BC x Htf** | **24** | **0.374** | **[0.225 - 0.56]** | **This study** |
|  | **BC x Hmm** | **24** | **0** | **[0 - 0.033]** | **This study** |

Figure S1. Map showing the geographic distribution and wing phenotype of *Heliconius melpomene malleti* and *H. timareta florencia.*

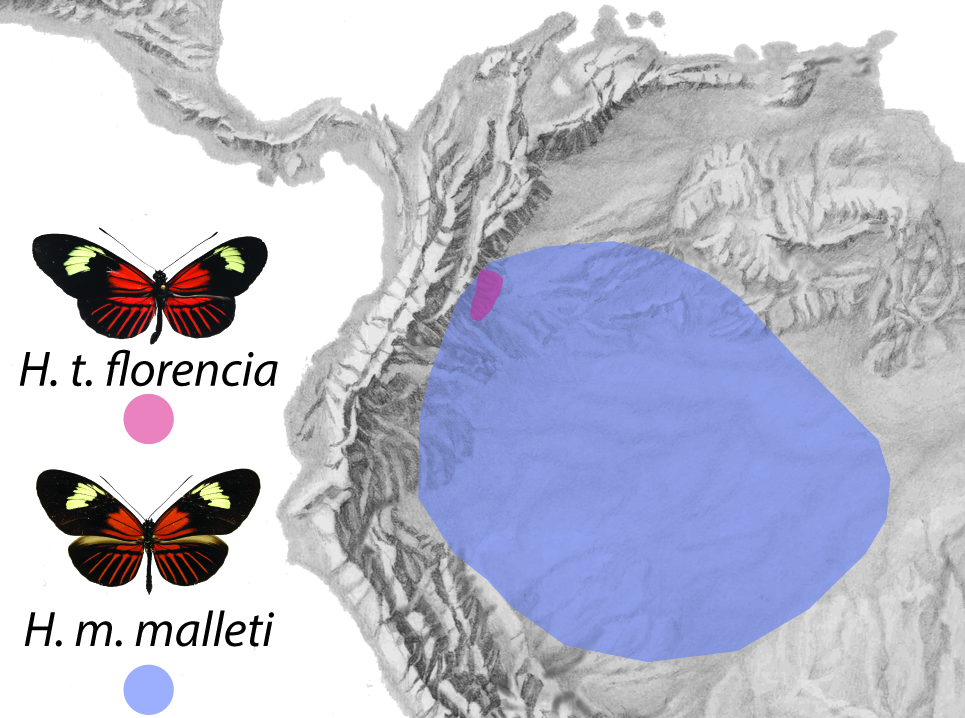

Figure S2. Location of landmarks on the forewing and hindwing of *H. melpomene malleti* and *H. timareta florencia,* used in the colour pattern analysis (**A**) and in the shape and size analysis (**B**). Landmarks are indicated by grey circles. Shapes deformation (**B)** and (**D)** represents the shape at minimum values for PC1 and PC2, respectively. Shapes deformation (**C)** and (**E**) represents maximum values for PC1 and PC2, respectively. Landmarks indicated (LM3, LM4 and LM15) are the one that varies the most.

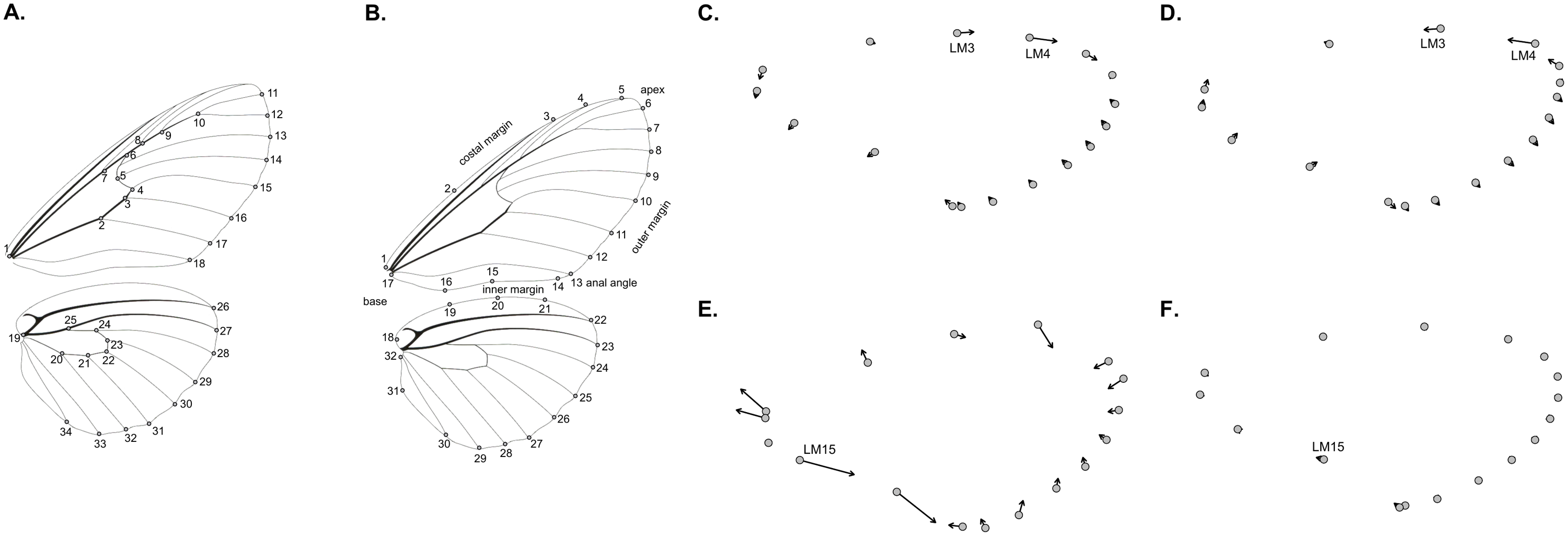

Figure S3. Species wing size and shape. Forewing (**A**) and hindwing (**B**) size variation. Density plots showing the variation in the shape of the forewing (**C**) and the hindwing (**D**). A total of 43 *H. m. malleti* and 45 *H. t. florencia* were analysed.

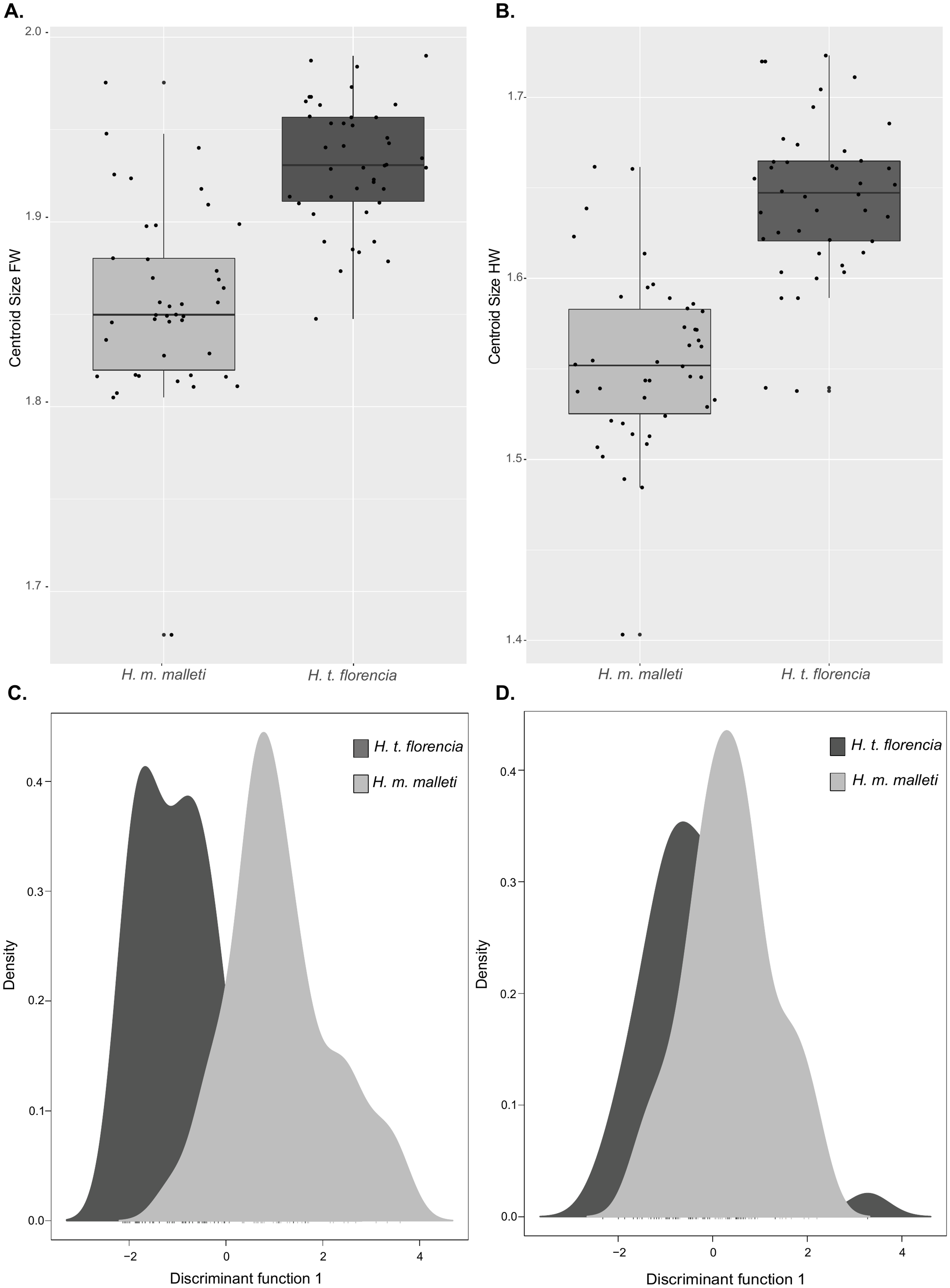

Figure S4. Shape variation of forewings of *H. melpomene malleti* (light grey) and *H. timareta florencia* (dark grey) (**A**). Panels (**B**) and (**C**) indicate PC1 and PC2 loadings respectively.

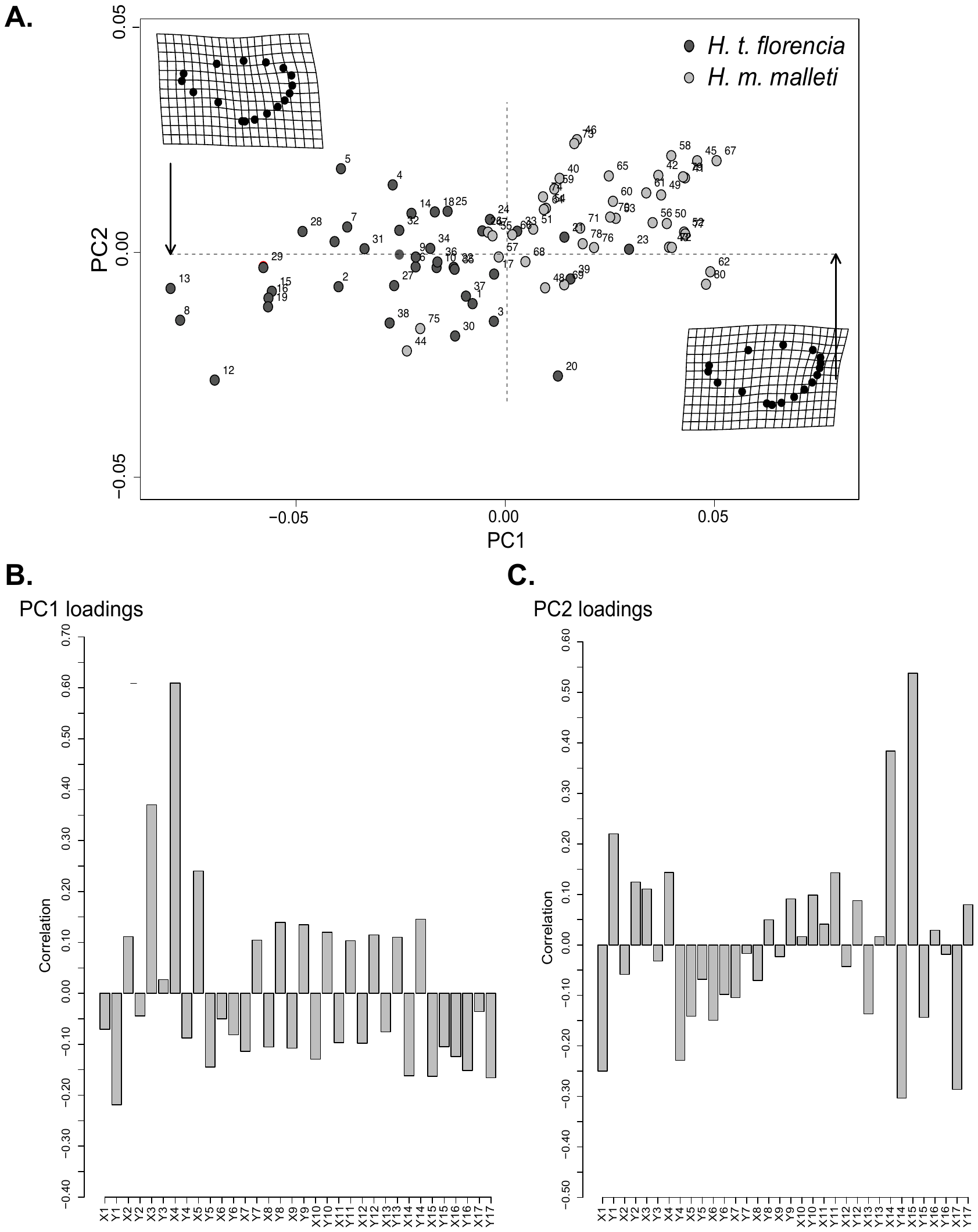

Figure S5. Colour pattern comparison between the two species. (**A**) Yellow patch on the dorsal forewing; (**B**) Dennis on the dorsal forewing; (**C**) Ray on the dorsal hindwing; (**D**) Yellow patch on the ventral forewing; (**E**). Dennis on the ventral forewing and (**F**) ray on the ventral hindwing. Yellow circle represents individuals of *H. t. florencia* and red triangles represents individuals of *H. m. malleti*. For each individual (n=88) only those wings in good condition were used (Table S1). Predicted values scale indicates the expression of colour pattern, where positive values correspond to higher presence (red) and negative values (blue) to absence of the colour pattern. PC1 explains variation in wing elements size and PC2 explain variation in wing elements shape.

### Figure S6. Amount (ng) of compounds remained in the wings of perfumed males with the heterospecific hexanic extract before evaporation.

We dissected the wings after 1 minute, 30 minutes and 60 minutes after spreading the heterospecific hexanic extract. (**A**) Octadecanal; (**B**) Heneicosane; (**C**) Syringaldehide; (**D**) Z-11-eicosanal. Blue line represents the pattern present in *H. m. malleti* and red line represents *H. t. florencia*.

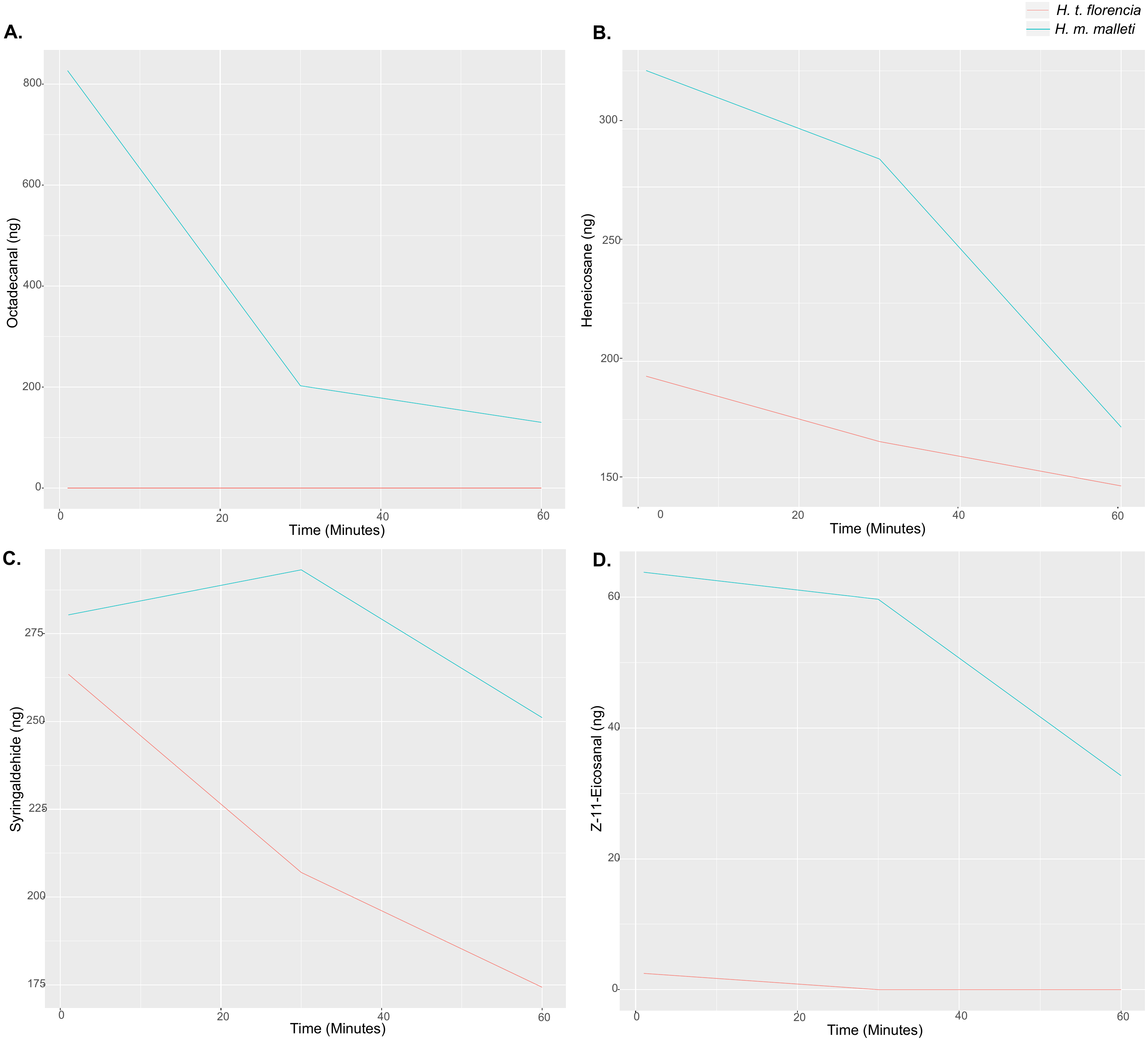

Figure S7. Female behavioural responses towards conspecific males “perfumed” with a hexane extract from five males either of *H. m. malleti* or *H. t. florencia*. Behaviours are classified as Acceptance (A) or Rejection (R). Control males are represented in red (left) and treatment males in blue (right). Means are marked with a black square and boxplots mark the inter-quartile ranges. Asterisk (*) next to the species’ name is indicative of statistically significance (α=0.01) according to GLMM.

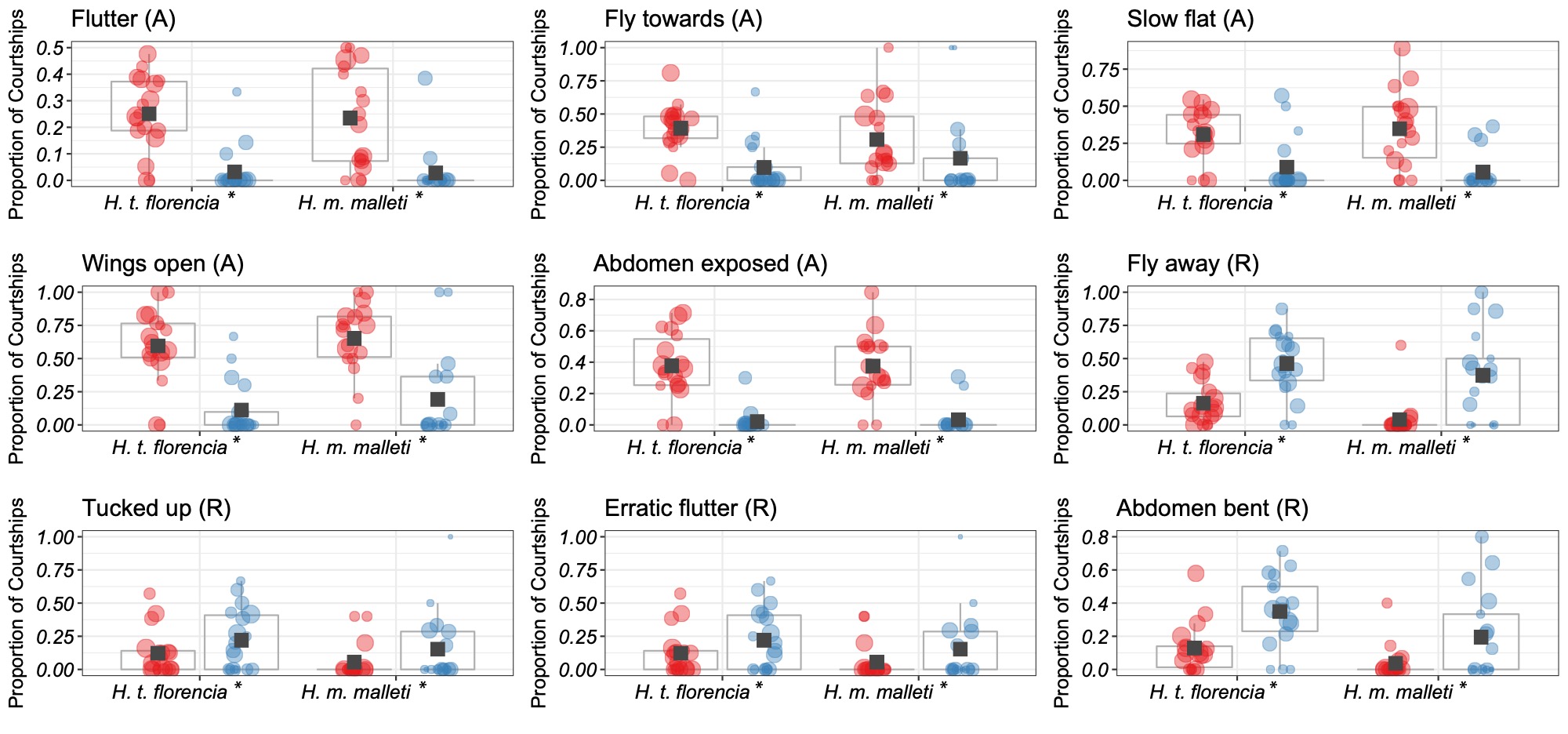

Figure S8. Mate choice triads testing behavioural responses in F_1_ and backcrosses (BC) females (**A**). Number of mating obtained is indicated above each bar. *H. t. florencia* males are represented in light grey and *H. m. malleti* males are represented in dark grey. (**B**) Proportion of courtships that resulted in behavioural responses in F_1_ and BC females. Behaviours recorded were acceptance (A) or rejection (R) towards males of *H. m. malleti* (red, left) and *H. t. florencia* (blue, right). Means are marked with a black square and boxplots mark the inter-quartile ranges. Size of datapoint is proportional to the number of courtships by that male. Asterisk * next to the female (F_1_/BC) is indicative of statistically significance (α=0.01) according to GLMM.

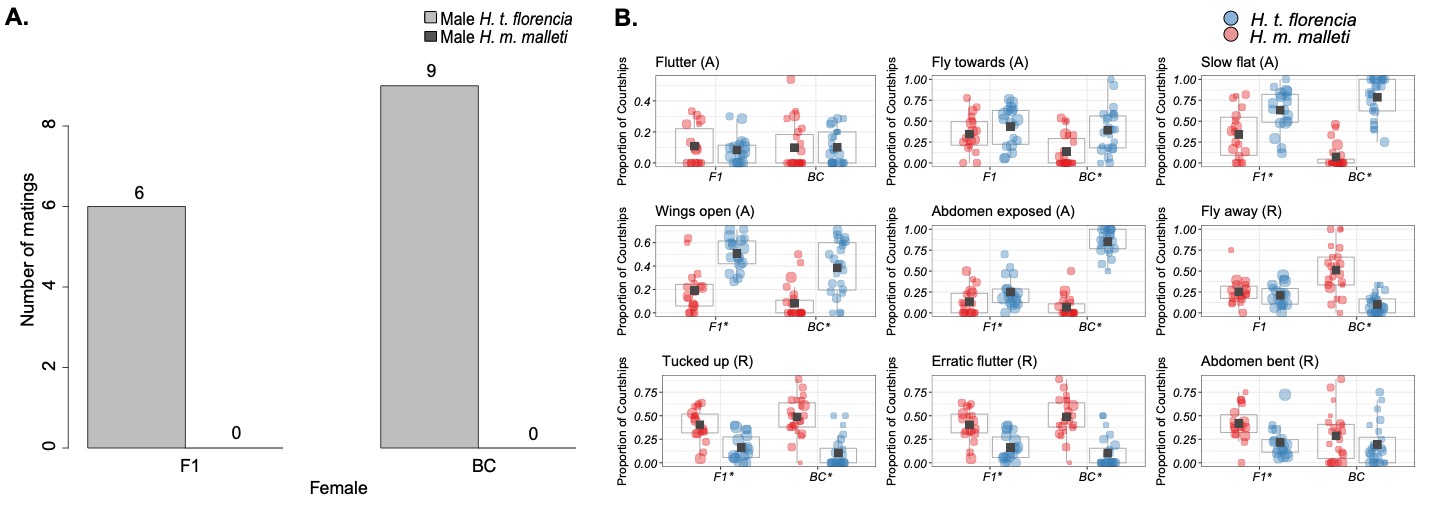

### Figure S9. Species differences in male androconial extracts.

Chromatogram of extract of androconial region from (**A**) *H. timareta florencia* and (**B**) *H. melpomene malleti*. IS, internal standard (2-tetradecylacetate); 1, dihydroactinidiolide; 2, unknown; 3, syringaldehide; 4, henicosane; 5, octadecanal.

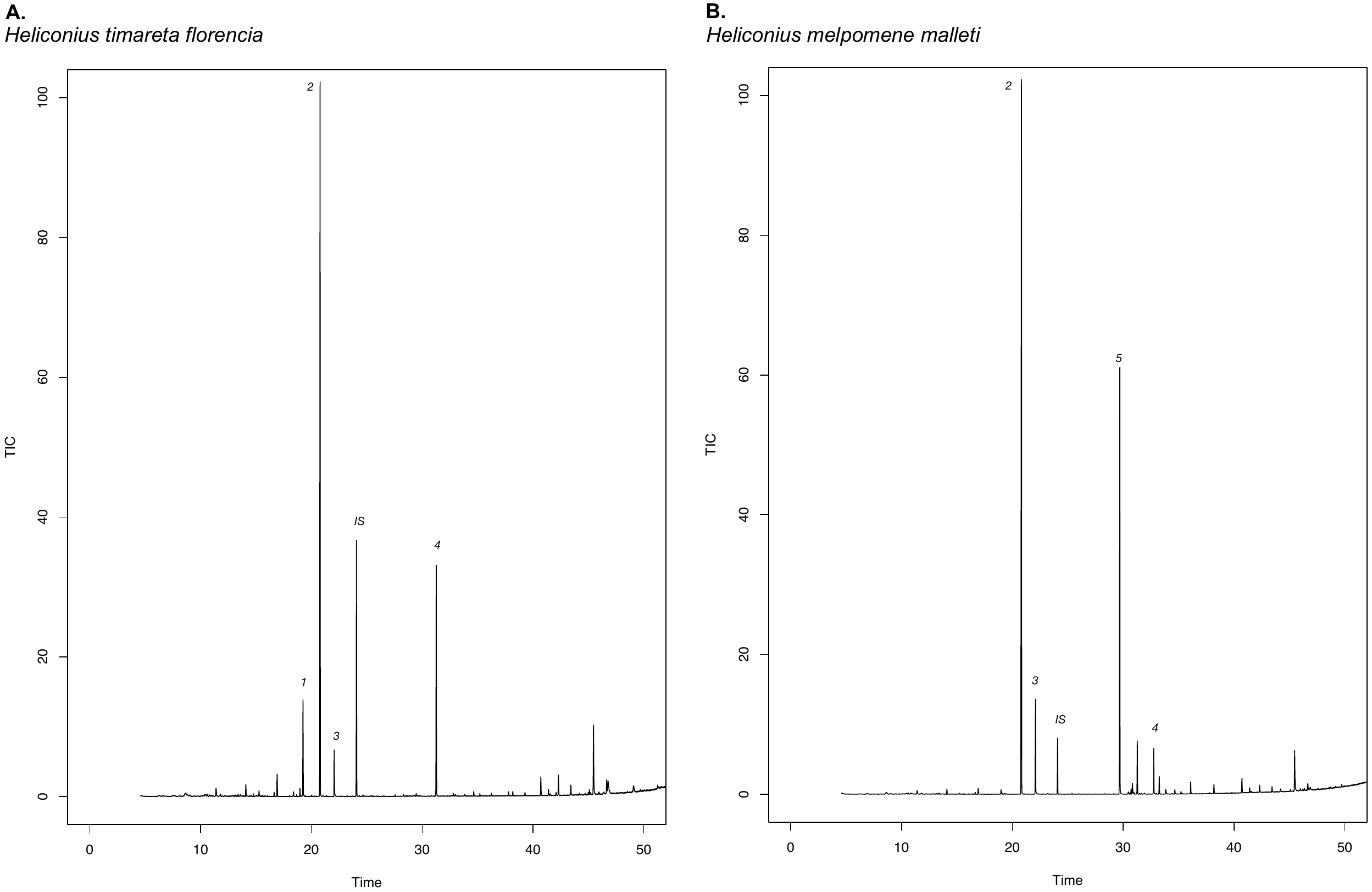

Figure S10. Cluster analysis based on Euclidian distance of compound composition in the wing androconia of males of *H. m. malleti*, *H. t. florencia*, F_1_ and BC. In red the most abundant compounds in the wing androconia.

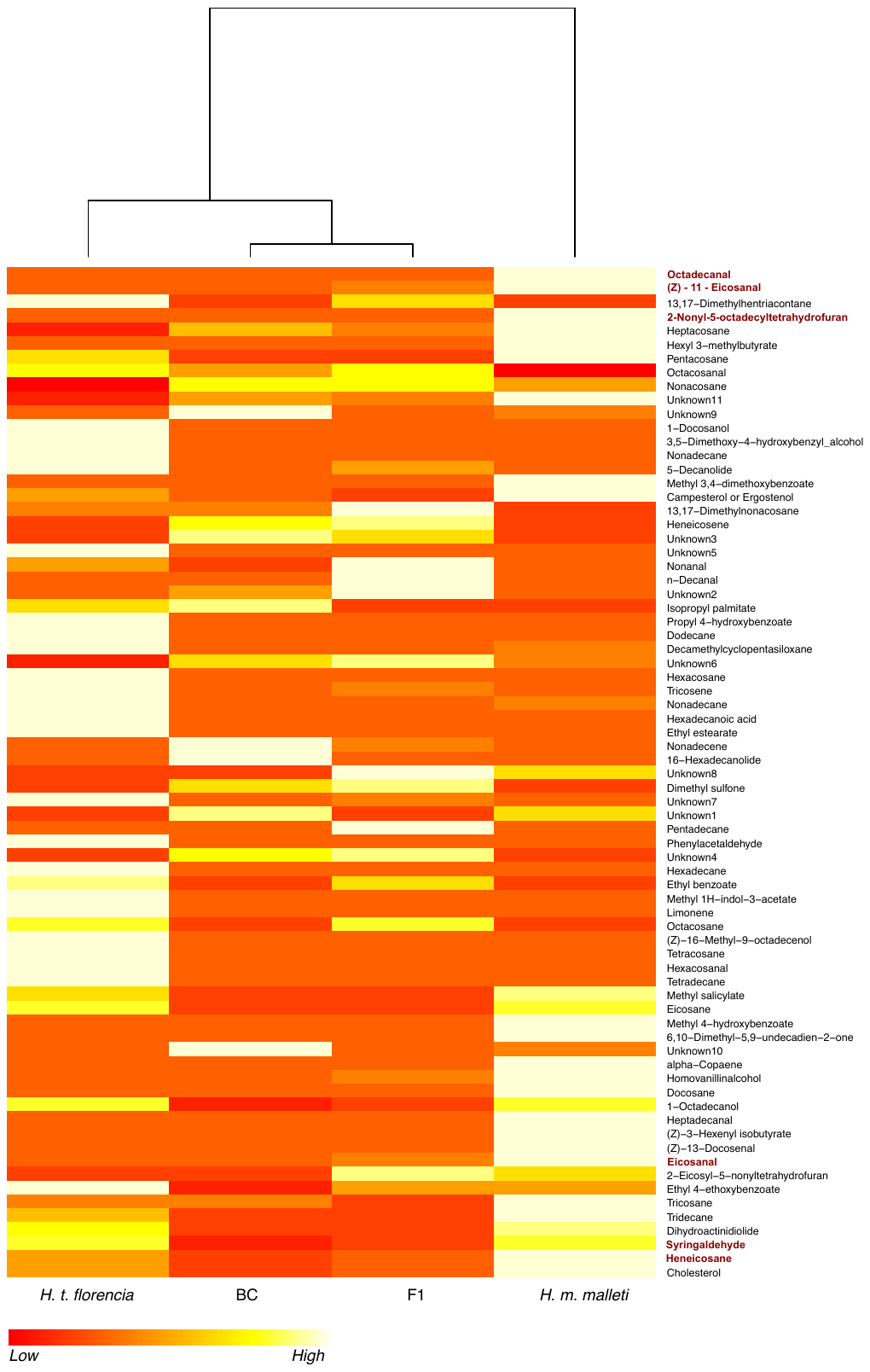

Figure S11. Differences in androconial extracts of hybrid F_1_ and backcrosses males. Chromatograms of extract of androconial region from different (**A**) F_1_ and (**B**) backcrosses individuals. IS, internal standard (2-tetradecylacetate); 1, dihydroactinidiolide; 2, unknown; 3, syringaldehide; 4, henicosane; 5, octadecanal.

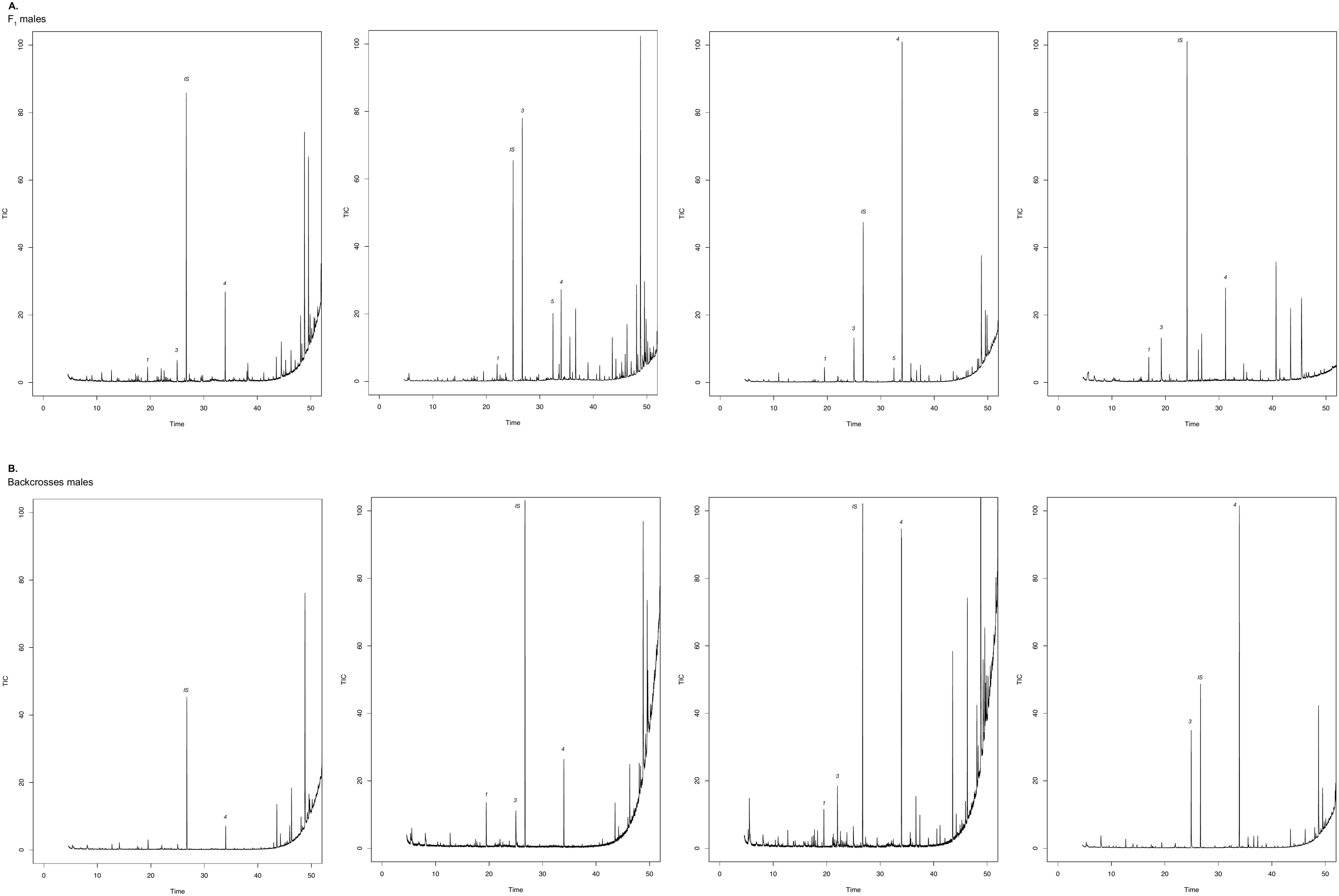

Figure S12. Species differences in male abdominal gland bouquet. Chromatogram of extract of abdominal gland from (**A**) *H. timareta florencia* and (**B**) *H. melpomene malleti*. IS, internal standard (2-tetradecylacetate); 1, eicosane; 2, Z-9-C18,11olide; 3, heneicosane; 4, ethyl oleate; 5, isopropyl oleate; 6, isopropyl octadecanoate; 7, butyl oleate; 8, β-ocimene; 9, henicosene.

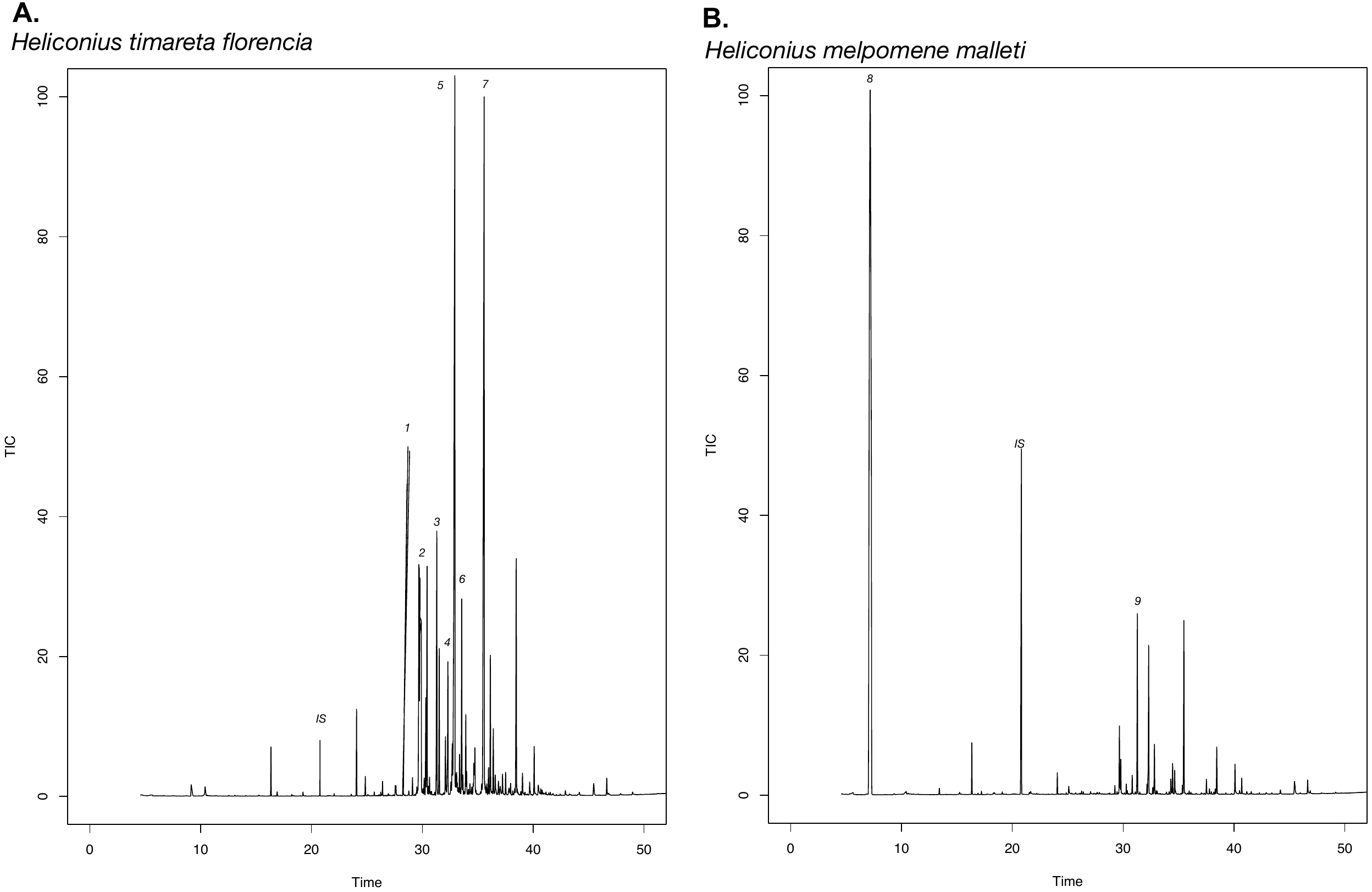

Figure S13. Cluster analysis based on Euclidian distance of compound composition in the wing androconia of males of *H. m. malleti*, H*. t. florencia*, F_1_ and BC. In red the most abundant compounds in the wing androconia.

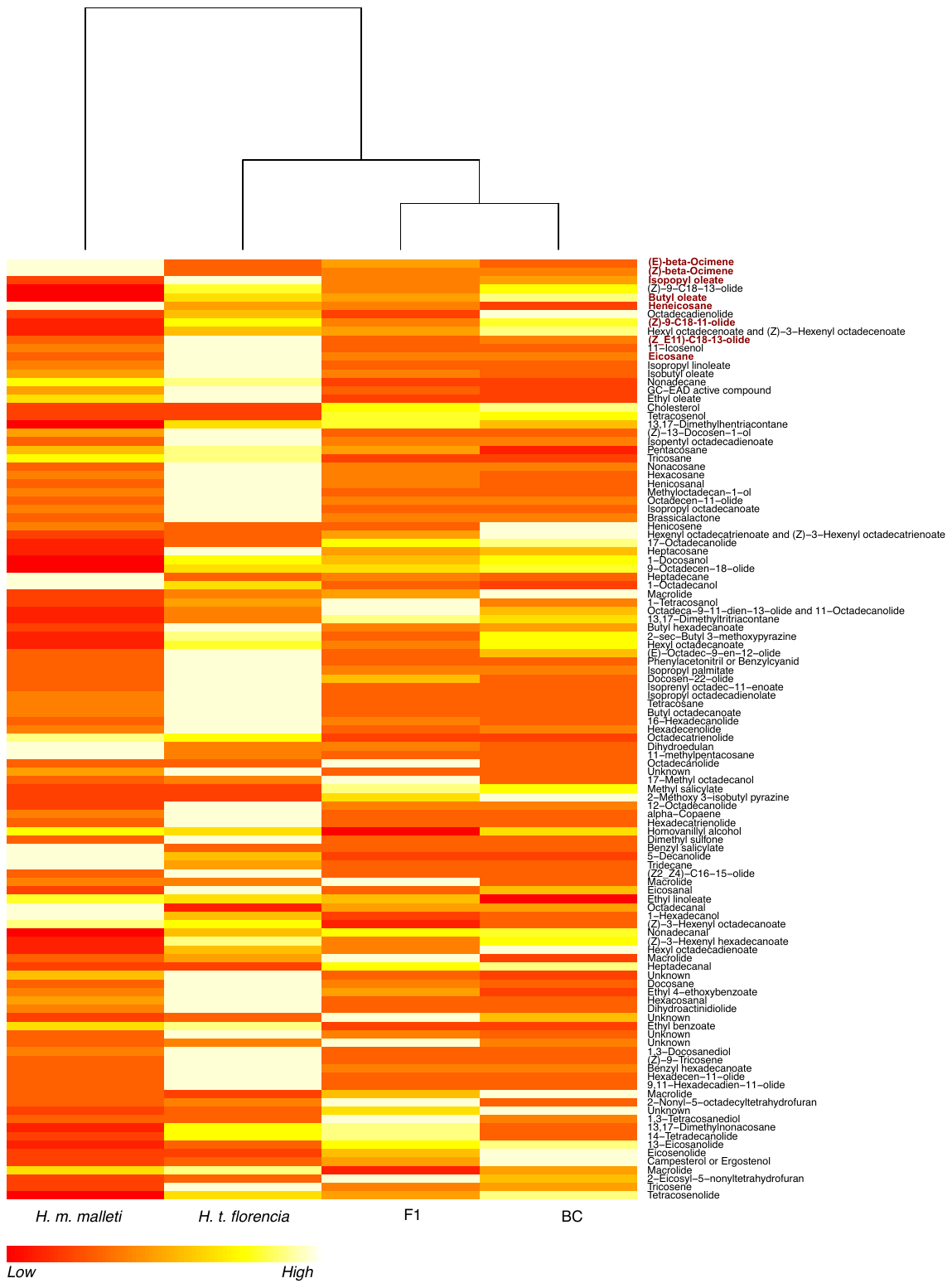

### Figure S14. Differences in the abdominal gland bouquet of hybrid F_1_ and backcrosses males.

Chromatograms of extract of abdominal gland from (**A**) F_1_ and (**B**) backcrosses individuals. IS, internal standard (2-tetradecylacetate); 1, eicosane; 2, Z-9-C18,11olide; 3, heneicosane; 4, ethyl oleate; 5, isopropyl oleate; 6, isopropyl octadecanoate; 7, butyl oleate; 8, β-ocimene; 9, henicosene.

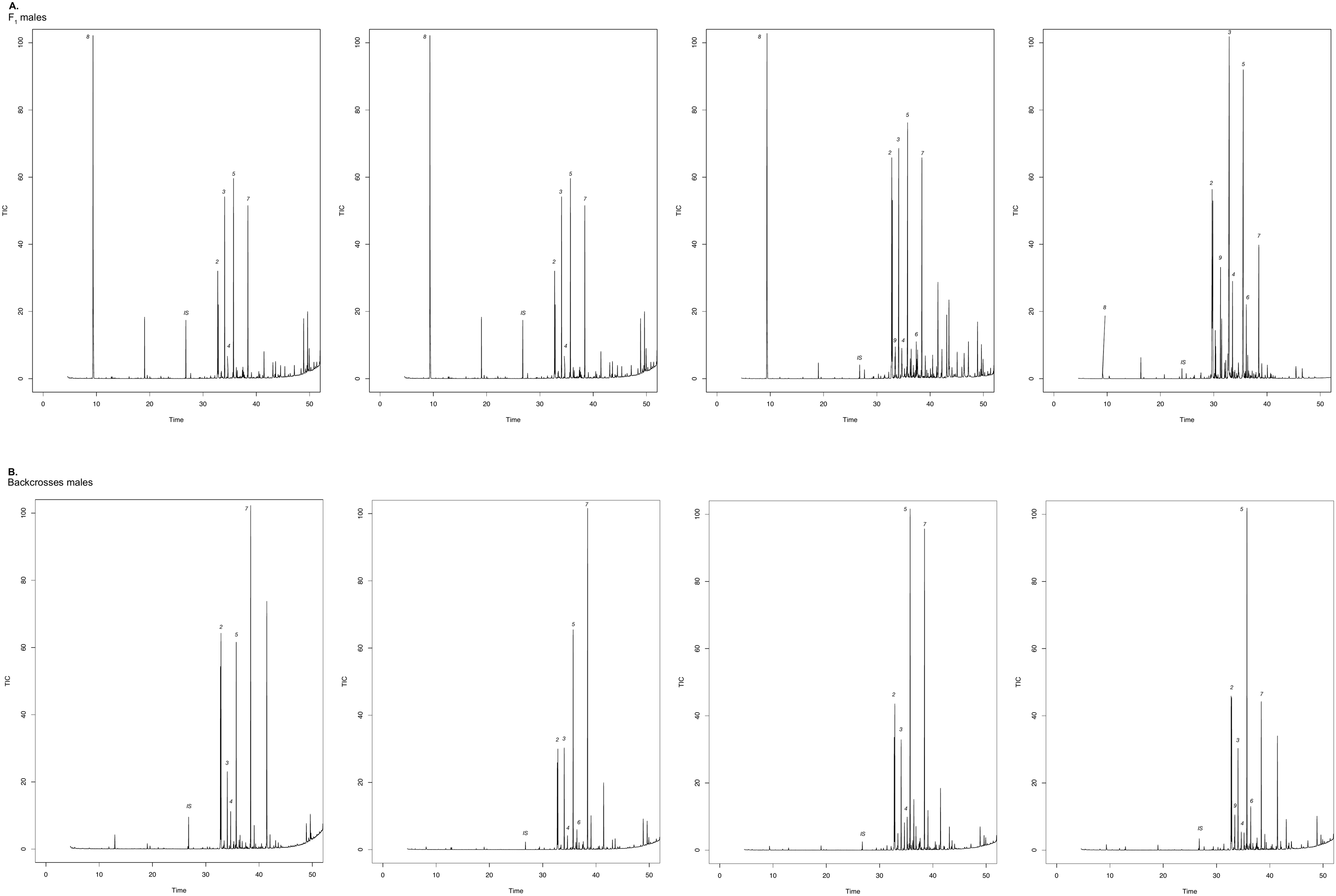
